# Production and characterization of virus-free, CRISPR-CAR T cells capable of inducing solid tumor regression

**DOI:** 10.1101/2021.08.06.455489

**Authors:** Katherine P. Mueller, Nicole J. Piscopo, Matthew H. Forsberg, Louise A. Saraspe, Amritava Das, Brittany Russell, Madeline Smerchansky, Lei Shi, Keerthana Shankar, Adeela Ali, Cicera R. Lazzarotto, Shengdar Q. Tsai, Christian M. Capitini, Krishanu Saha

## Abstract

**Background:** Chimeric antigen receptor (CAR) T cells traditionally harbor viral vector-based sequences that encode the CAR transgene in the genome. These T cell products have yet to show consistent anti-tumor activity in patients with solid tumors. Further, viral vector manufacturing is resource intensive, suffers from batch-to-batch variability, and includes several animal components, adding regulatory and supply chain pressures.

**Methods:** Anti-GD2 CAR T cells were generated using CRISPR/Cas9 within nine days using recombinant Cas9 protein and nucleic acids, without any viral vectors or animal components. The CAR was specifically targeted to the T Cell Receptor Alpha Constant gene (*TRAC*). T cell products were characterized at the level of the genome, transcriptome, proteome, and secretome using CHANGE-seq, scRNA-seq, spectral cytometry, and ELISA assays. Functionality was evaluated *in vivo* in an NSG xenograft neuroblastoma model.

**Results:** In comparison to traditional retroviral CAR T cells, virus-free CRISPR CAR (VFC-CAR) T cells exhibit *TRAC*-targeted genomic integration of the CAR transgene, elevation of transcriptional and protein characteristics associated with a memory phenotype, and low tonic signaling prior to infusion arising in part from the the knockout of the TCR. Upon exposure to the GD2 target antigen, anti-GD2 VFC-CAR T cells exhibited specific cytotoxicity against GD2+ cells *in vitro* and induced solid tumor regression *in vivo*, with robust homing, persistence, and low exhaustion against a human neuroblastoma xenograft model.

**Conclusions:** This proof-of-principle study leveraging virus-free genome editing technology could enable flexible manufacturing of clinically relevant, high-quality CAR T cells to treat cancers, including solid tumors.

## Text

Chimeric antigen receptor (CAR) T cell therapy is rapidly transforming the treatment of many cancers, with five products already approved by the Food and Drug Administration for some hematologic malignancies. However, solid tumors have presented a difficult challenge for the CAR T field, as clinical trials to date have yielded little to no responses and no improvement in survival^1,2^ due in part to poor T cell potency and/or persistence within patients^3^. New CAR T products, manufactured for high potency within solid tumors, are critically needed^4–6^.

CAR T cells are traditionally manufactured using lentiviruses or *γ*-retroviruses^7,8^ which confer high-efficiency editing; however, viral transduction methods broadly integrate their nucleic acid payloads into the host genome, risking insertional mutagenesis^7,9^. In addition, poorly specified integration of a CAR transgene can lead to heterogeneous and unpredictable CAR expression. Good manufacturing practice (GMP)-grade viral vectors and associated quality testing are also expensive and constitute a major supply chain bottleneck for the field^10^. Nonviral methods of CAR transduction include transposon-mediated integration^11^ and transient mRNA delivery through electroporation^12^. Like lenti- and retroviruses, transposons also integrate the CAR broadly throughout the genome, while mRNA-mediated delivery results in only transient CAR expression over a period of days, which can be problematic for achieving sustained remission^13^. Therefore, standard nonviral delivery methods present considerable challenges for precise and durable CAR gene transfer.

Recent strategies employing viral vectors and CRISPR/Cas9 genome editing^14–16^ have targeted the CAR transgene to a single genomic locus to reduce the risks of insertional mutagenesis and transgene silencing. Eyquem and colleagues inserted an anti-CD19 CAR into exon 1 of the T cell receptor alpha chain (*TRAC*), disrupting expression of the T cell receptor (TCR) while also driving CAR expression from the endogenous *TRAC* promoter^17^. These T cells, engineered through electroporation of Cas9 mRNA followed by delivery of a homology-directed repair (HDR) template within a recombinant adeno-associated viral (AAV) vector, were potent, retained a memory phenotype, and showed less exhaustion relative to conventional *γ*-retroviral products. An AAV-mediated approach has also been combined with TALEN technology to engineer *TRAC*-targeted CAR T cells, with comparable effects on T cell phenotype^18^. These phenotypes correlate with improved outcomes for patients with hematological malignancies^19–26^. Strategies that lead to memory CAR T cell generation with lower exhaustion and terminal differentiation phenotypes have been hypothesized to be beneficial in treating solid tumors^27^. The use of AAVs to deliver the HDR template needed for CRISPR-mediated transgene insertion^17,28^ is however limited by supply chain challenges associated with viral vector production^10^. Additionally, vector integration into the genome with AAVs can occur when used in conjunction with Cas9^28^, and cellular response to the introduction of viral elements could affect T cell phenotypes. Therefore, alternate strategies for precise CAR transgene insertion that avoid viral vectors entirely could yield new opportunities to flexibly manufacture CAR T cell immunotherapies with desirable phenotypes.

Completely virus-free CRISPR/Cas9-mediated gene transfer with transgenic TCR and anti-CD19 CARs has recently been demonstrated to be functional against some cancers^29–31^, but not for solid tumors. Here, we build upon these virus-free methods^29,32^ to integrate a 3.4 kb third-generation **anti-disialoganglioside** (GD2) CAR transgene^33^ at the human *TRAC* locus to report a completely **virus-free CRISPR CAR (VFC-CAR)** T cell product featuring precise genomic integration of a CAR that has been validated in an *in vivo* solid tumor model. VFC-CAR T cells exhibit more transcriptional and protein expression characteristics associated with a memory-like phenotype relative to conventional ***γ*-retroviral (RV)**-CAR T cells. These VFC-CAR T cells also show evidence of decreased TCR and CAR signaling prior to antigen exposure and comparable potency relative to conventional viral CAR T cells against GD2+ neuroblastoma *in vivo*.

## Results

### VFC-CAR T cells can be efficiently manufactured with low CAR expression heterogeneity

To avoid using HDR donor templates within viral vectors, we first cloned a third generation GD2-targeting CAR sequence^33^ into a plasmid containing homology arms flanking the desired cut site at the start of the first exon of the *TRAC* locus (**figure 1A**). The same third generation GD2-targeting CAR sequence was used to generate RV-CAR T cells as a comparison throughout this study (**figure 1B**). We next generated double-stranded DNA (dsDNA) HDR templates via PCR amplification off the plasmid and performed a two-step purification process to purify and concentrate the templates. Building on prior established protocols^29^, we performed two sequential purifications on the PCR amplicons to produce a highly-concentrated dsDNA HDR template. Primary human T cells from healthy donors were electroporated with the HDR templates and Cas9 ribonucleoproteins (RNPs) targeting the human *TRAC* locus. Cells were allowed to recover for 24 hours at high density in round-bottom 96-well plates. Next, the cells were cultured in xeno-free media and assayed on days 7 and 9 post-isolation to produce VFC-CAR T cell products. We also include a **virus-free CRISPR control (VFC-Ctrl)** condition in which cells harbor the same disruption of the *TRAC* locus, but with a signaling-inert mCherry fluorescent protein inserted in place of the CAR (**figure 1B**).

**Figure 1.**
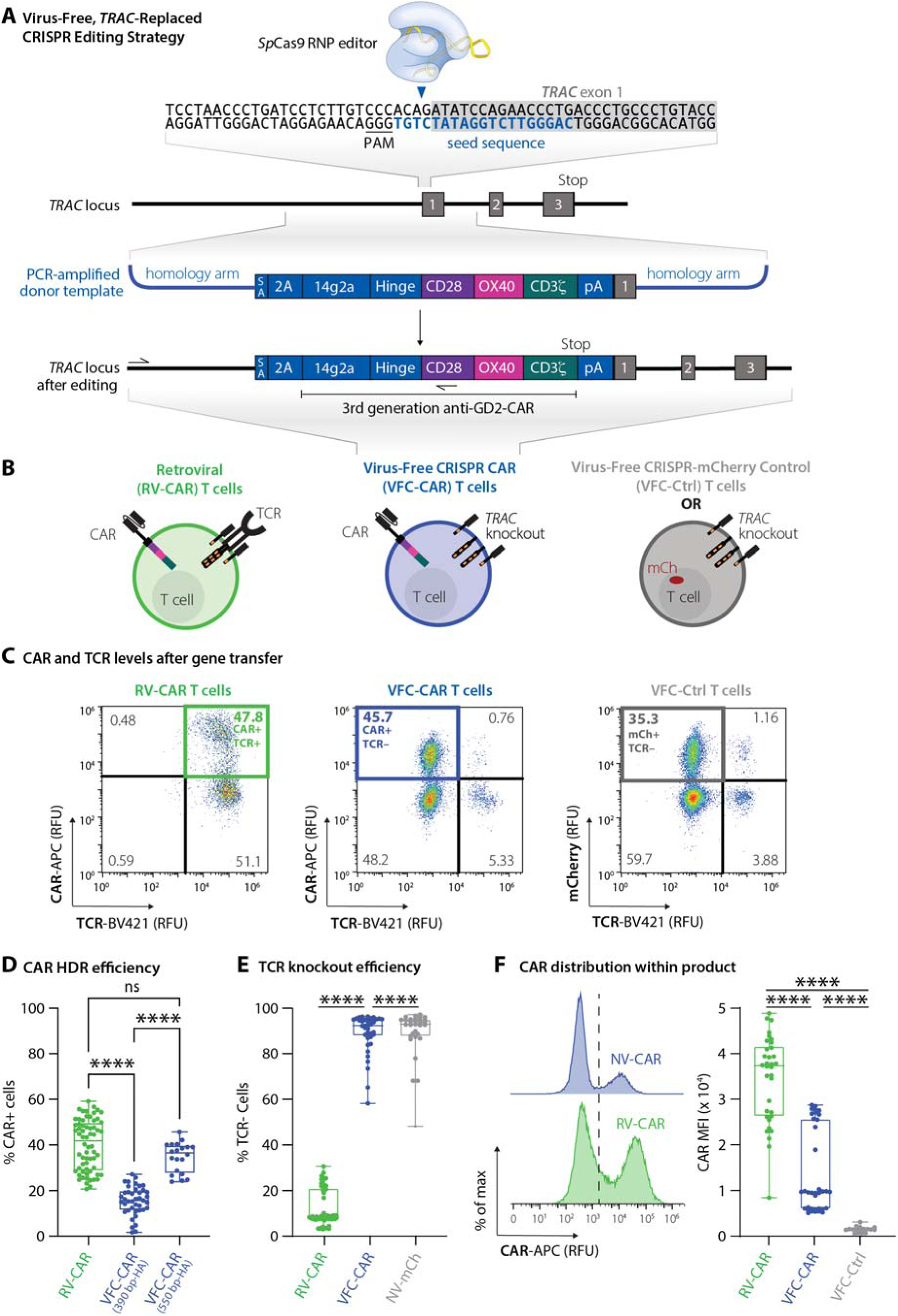
Virus-free CRISPR (VFC)-CAR T cells are efficiently generated in one step by replacing the T cell receptor with the CAR. (A) Schematic showing the CAR genetic construct and virus-free strategy to insert the CAR into the first exon (grey box) of the human *TRAC* gene. No viral components are necessary, and the CRISPR-Cas9 ribonucleoprotein is delivered transiently via electroporation. The seed sequence of the gRNA is in blue and the protospacer adjacent motif (PAM) for SpCas9 is underlined. 14g2a: single chain variable fragment clone targeting GD2; SA: splice acceptor, 2A: self-cleaving peptide, pA: rabbit ß-globin polyA terminator. Arrows indicate positions of primers for in-out PCR assay shown in figure 2. (B) Schematic of T cell products used in this study with receptors and expressed transgenes. VFC-CAR, virus-free CRISPR CAR T cell product generated by electroporation. RV-CAR, donor-matched CAR T cell product generated by retroviral transduction with the same third generation anti-GD2 CAR shown in A; VFC-Ctrl, donor-matched control T cell product manufactured as in A but with an mCherry fluorescent protein instead of a CAR. (C) Representative density flow cytometry plots for transgene and TCR surface protein levels on the manufactured cell products. Y-axis shows CAR or mCherry levels and X-axis shows TCR levels on day 7 post-isolation (day 5 post-electroporation for VFC-CAR and VFC-Ctrl, and day 4 post-transfection for control RV-CAR). Thick colored boxes delineate cell populations selected for downstream analysis. (D) Boxplots show the percentage of CAR positive cells from gene editing for VFC-CAR cells and from retroviral transduction for RV-CAR cells in each sample The first VFC-CAR product featured homology arms (HA) of 383 (left) and 391 (right) bp, respectively. The homology arms on the second VFC-CAR product were extended to 588 (left) and 499 (right) bp, respectively. (E) Boxplots show the percentage of TCR negative cells from gene editing in VFC-CAR cells and in RV-CAR cells. RV-CAR TCR negativity likely results from endogenous repression of the TCR. (F) Mean fluorescence intensity (MFI) values for the CAR expression levels with associated histograms. Boxplots show the percentage of CAR positive cells in each sample. * indicates p≤0.05; ** indicates p≤0.01; *** indicates p≤0.001; **** indicates p≤0.0001.

We profiled each cell product for viability and yield at various points throughout the manufacturing process. The viability of VFC-CAR and RV-CAR T cells were both high (>80%) by the end of manufacturing (**online supplemental figure S1A**). Cell proliferation and growth over nine days were robust for both groups (**online supplemental figure S1A**). We assessed gene editing at multiple points post-isolation and achieved higher levels of CAR integration when cells were edited at 48 hours after CD3/CD28/CD2 stimulation (**online supplemental figure S1B**). Using these templates, we achieved consistently high genome editing across over 4 donors, with an average of 15% knockin efficiency. We improved targeting efficiency further by using alternate primer pairs in our PCR strategy, which increase the length of the homology arms from ∼390 bp to ∼550 bp on either side of the CAR. This product, while larger in size (3.4 kb), demonstrated up to 45% knockin efficiency, with an average of 34% CAR+ and TCR-cells, as measured by flow cytometry (**figure 1C, D**). Within the VFC-CAR samples, the TCR was consistently knocked out in >90% of T cells (**figure 1E**). The mean fluorescence intensity (MFI) of CAR expression was significantly elevated and showed greater range (∼1.6 fold; **figure 1F**) in the RV-CAR samples in comparison to the VFC-CAR samples, indicating decreased CAR expression heterogeneity within the VFC-CAR product and consistent with prior findings with AAV-CRISPR-CAR T cells^4^.

### Genomic analysis indicates specific targeting of the CAR transgene to the *TRAC* locus

After confirming robust CAR protein expression, we performed genomic analysis to measure the on-target specificity of the gene edit. Proper genomic integration of the CAR was confirmed via an “in-out” PCR amplification assay^34^ on the genomic DNA extracted from the manufactured cell products with primers specific to the *TRAC* locus and the transgene (**figure 2A**). Next-generation sequencing of genomic DNA to profile *TRAC* alleles in the cell products without an integrated transgene confirmed high rates of genomic disruption at the *TRAC* locus for these residual alleles, with 93.06% indels for the VFC-CAR and VFC-Ctrl samples. Altogether, the combined genomic integration of the CAR or mCherry transgene and indels at the *TRAC* locus resulted in concomitant loss of TCR protein on the T cell surface in sample-matched assays (**figure 2B, C**). Genome-wide, off-target activity for our editing strategy was assayed by CHANGE-seq^35^. The top modified genomic site was identified to be the intended on-target site (**figure 2D, E**) with a rapid drop-off for off-target modifications elsewhere in the genome. The CHANGE-seq specificity ratio of our *TRAC* editing strategy is above average (0.056; 57^th^ percentile) when compared to published editing strategies previously profiled by CHANGE-seq^35^.

**Figure 2.**
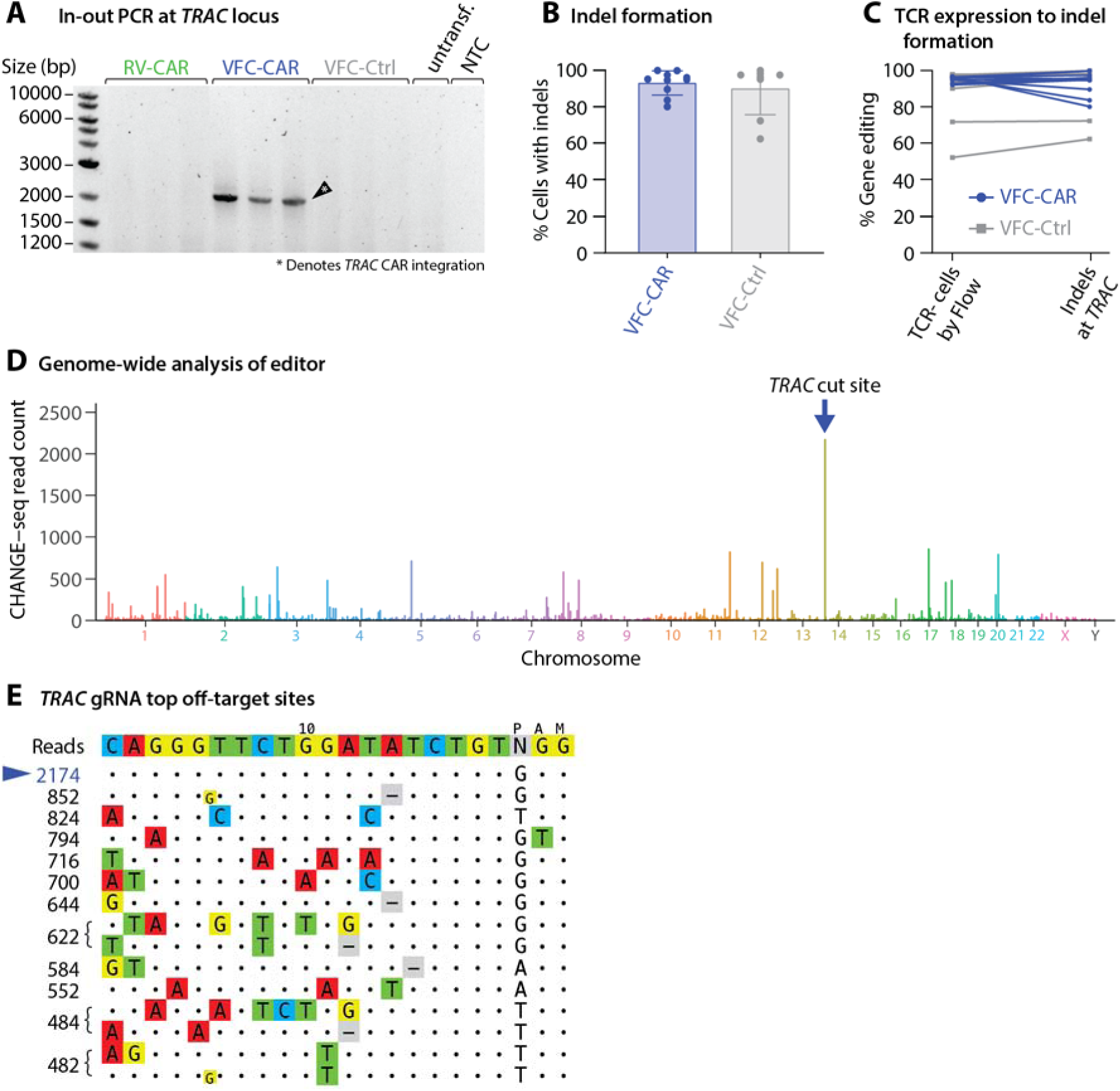
VFC-CAR T cells are efficiently and specifically edited at the *TRAC* gene. (A) In-out PCR indicates proper on-target genomic integration of the CAR transgene in VFC-CAR cells. Primer locations are shown in figure 1A by arrows upstream of the left homology arm and within the CD28 sequence of the CAR. Untransf., untransfected donor-matched T cells; NTC, no template control. (B) Percent of cells with indels at the *TRAC* gene in both VFC-CAR and VFC-Ctrl conditions. VFC-CAR (blue) N=10; VFC-Ctrl (grey) N=8, both for one donor. (C) Level of TCR editing in VFC-CAR and VFC-Ctrl T cells measured by both flow cytometry (left) and deep sequencing of genomic DNA (presence of insertions and deletions, indels, at the *TRAC* locus, right). VFC-CAR (blue) N=10, VFC-Ctrl (grey) N=8. (D) Manhattan plot of CHANGE-seq-detected on- and off-target sites organized by chromosomal position with bar heights representing CHANGE-seq read count. The on-target site is indicated with the blue arrow. (E) Visualization of sites detected by CHANGE-seq. The intended target sequence is shown in the top line. Cleaved sites (on- and off-target) are shown below and are ordered top to bottom by CHANGE-seq read count, with mismatches to the intended target sequence indicated by colored nucleotides. Insertions are shown in smaller lettering between genomic positions, deletions are shown by (-). Output is truncated to top sites; additional sites are shown in online supplemental table S1.

### Cytokine profiling reveals high antigen-specific response for VFC-CAR T cells

After harvesting CAR T cells, we profiled secreted cytokines typically associated with a proinflammatory response. On day 9 of manufacturing prior to antigen exposure, RV-CAR T cells produced higher levels of IFNγ, TNFα, IL-2, IL-4, IL-10, IL-13, IL-6, IL-1β and IL-12p70, in comparison to both VFC-CAR and VFC-Ctrl T cells (**figure 3A**, individual replicates shown in **online supplemental figure S2A**). To determine cytokine production after antigen-induced stimulation, we performed a 24 hour co-culture between the engineered T cells and GD2+ CHLA20 neuroblastoma, then measured cytokines in the conditioned media. Interestingly, we found that in the presence of antigen stimulation, the previous trend had reversed: VFC-CAR T cells either matched or surpassed the level of cytokine production of the RV-CAR T cells (**figure 3A**). This result suggests that cytokine secretion after a single antigen stimulation is comparable in VFC-CAR and RV-CAR T cells, but basal secretion in the absence of antigen stimulation is decreased in VFC-CAR T cells.

**Figure 3.**
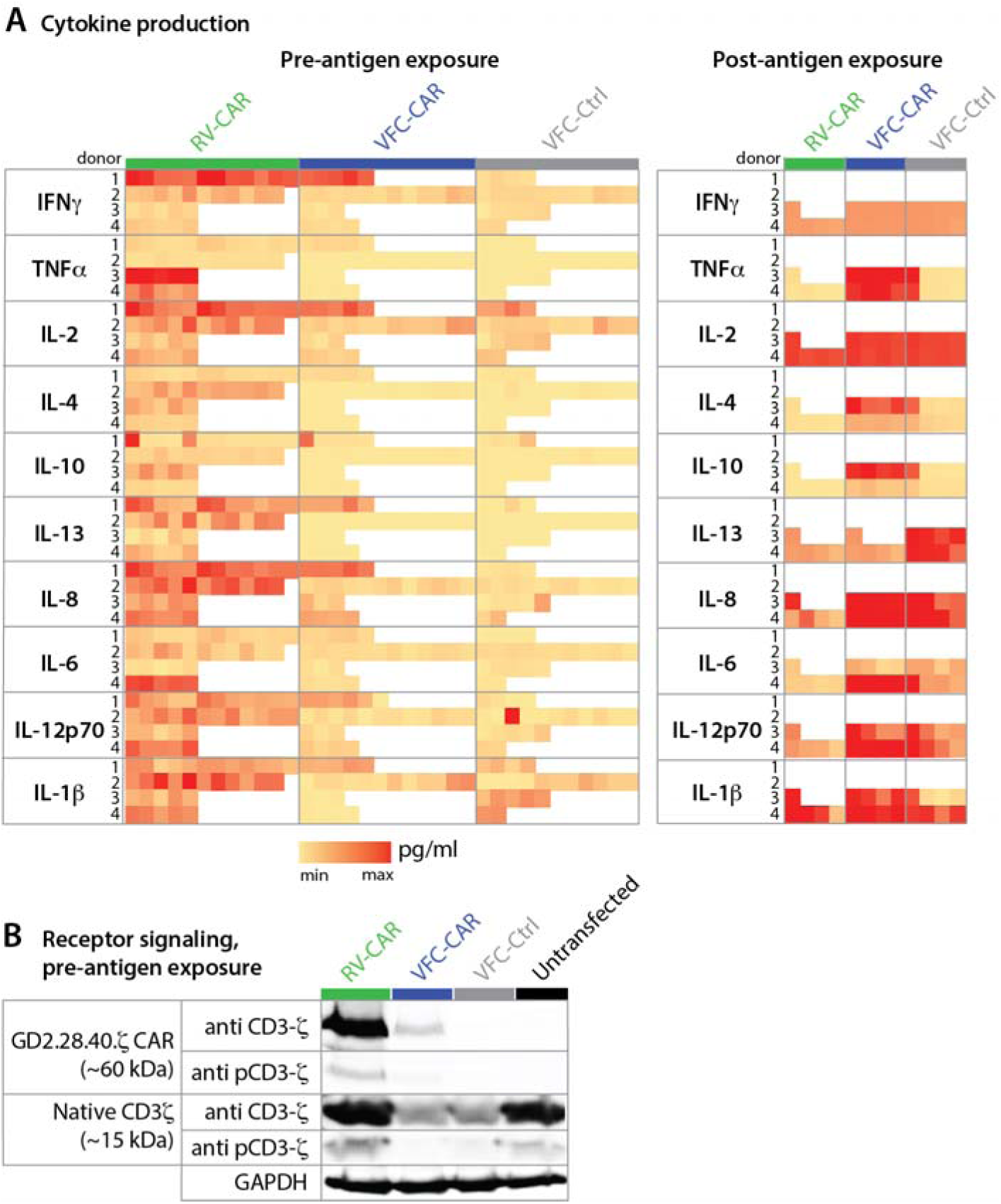
VFC-CAR T cells mount robust cytokine response upon exposure to cognate antigen, and decreased CAR and TCR-mediated signaling during manufacturing. (A, left) Cytokine production from conditioned media taken from T cell products at the end of manufacturing (pre-antigen exposure). Values are pooled from four donors. VFC-CAR (blue) N=24; RV-CAR (green) N=33; VFC-Ctrl (gray) N=22. (A, right) Cytokine production in conditioned media after a 24 hour co-culture of manufactured T cell products with the target antigen GD2 on CHLA20 neuroblastoma cells. Values are pooled from two donors. VFC-CAR (blue) N=8; RV-CAR (green) N=5; VFC-Ctrl (gray) N=8. The minimum and maximum for the color scale for each cytokine is as follows: IFN-y, 2000-30000 pg/mL. TNF-α, 10-1000 pg/mL. IL-2, 700-6000 pg/mL. IL-4, 0.2-20 pg/mL. IL-10, 0.5-150 pg/mL. IL-13, 300-1100 pg/mL. IL-8, 25-1800 pg/mL. IL-6, 10-400 pg/mL. IL-p70, 0.1-4 pg/mL. IL-1β, 0.2-100 pg/mL. Individual replicates are shown in **online supplemental figure S2.** (B) Western blot from cell lysates containing equivalent fractions of transgene+ cells (40% of each sample) and stained for CD3ζ, phosphorylated (p) CD3ζ, and GAPDH. CD3ζ domains from native CD3ζ and GD2.28.40.ζ were distinguished by molecular weight (15 and 60 kDa, respectively). N=1 donor.

### VFC-CAR T cells exhibit low basal TCR and CAR signaling during manufacturing

To test the possibility that variation in cytokine production prior to cognate antigen exposure resulted from varying levels of basal signaling from the CAR and/or TCR during *ex vivo* culture, we assayed CD3ζ phosphorylation from both native CD3ζ and the CD3ζ portion of the CAR via western blot (**figure 3B**). We found elevated protein levels of both CAR and TCR-associated CD3ζ in RV-CAR T cells relative to VFC-CAR T cells, potentially indicative of both a higher CAR copy number in RV products and an intact TCR-CD3ζ complex in the absence of *TRAC* knockout^36^. We also observed higher levels of CD3ζ phosphorylation in RV-CAR T cells from both CAR and TCR-associated protein, indicating elevated levels of basal signaling. In the absence of antigen exposure, elevated CAR/TCR signaling is likely present in the traditional RV-CAR T cells, a phenotype that has been associated with an increased propensity for terminal differentiation and exhaustion in some CAR T cell products, and which has been specifically identified in the context of CAR T cell products dependent on γ−retroviral vectors^37–39^. VFC-CAR and VFC-Ctrl cells both showed sharply decreased TCR-mediated CD3ζ signaling after *TRAC* knockout, and VFC-CAR T cells also showed minimal activity from CAR-associated CD3ζ. These results with our anti-GD2 CAR are consistent with prior findings of lower tonic signaling with an anti-CD19 CAR when CAR expression was driven by the endogenous *TRAC* promoter^17^. Both TCR and CAR-mediated basal signaling are diminished by our VFC strategy, in comparison to traditional RV products.

### VFC-CAR T cells exhibit elevated surface memory markers

To further explore the differential state of viral and virus-free CAR T cell proteomes, we performed an immunophenotyping panel using spectral cytometry, assaying for markers of T cell memory and differentiation state, activation, trafficking, exhaustion, and senescence (**online supplemental figure S3**). For all markers, we gated cells first by size and shape, then by viability, CD45 expression, and transgene expression to evaluate the multidimensional immunophenotypes within our products. We noted a dramatic decrease in CD3 expression in both VFC-CAR and VFC-Ctrl products relative to RV-CAR T cells, as expected following TCR knockout^36^. This finding corroborates the decrease in CD3ζ detected by western blotting (**figure 4A**). We next assessed expression of CD45RA and CD45RO, which are frequently used to distinguish naive/effector and memory subtypes^40^. Surprisingly, we found that a majority of cells expressed both markers, likely indicating a transitional cell state^41^; however, significantly more VFC-CAR and VFC-Ctrl T cells expressed the memory-associated CD45RO marker at high levels relative to RV-CAR T cells, indicating active formation of a central memory (T_cm_) or effector memory (T_eff_) phenotype (**figure 4B**). There was a skew toward high levels of CD62L, another memory-associated protein, in VFC products relative to viral products; this is consistent with phenotypes observed for *TRAC*-knockout CD19 CAR T cells^17^ (**figure 4B, online supplemental figure S4A**). The vast majority of cells in all groups expressed CD95, indicating that the cells have differentiated beyond a naive phenotype, as expected after activation by a CD2/CD3/CD28 tetrameric antibody in the culture media^42^ (**online supplemental figure S4A**). We noted that viral and VFC-CAR cells expressed comparable levels of the memory-associated protein CCR7, and significantly more than VFC-Ctrl cells.

**Figure 4.**
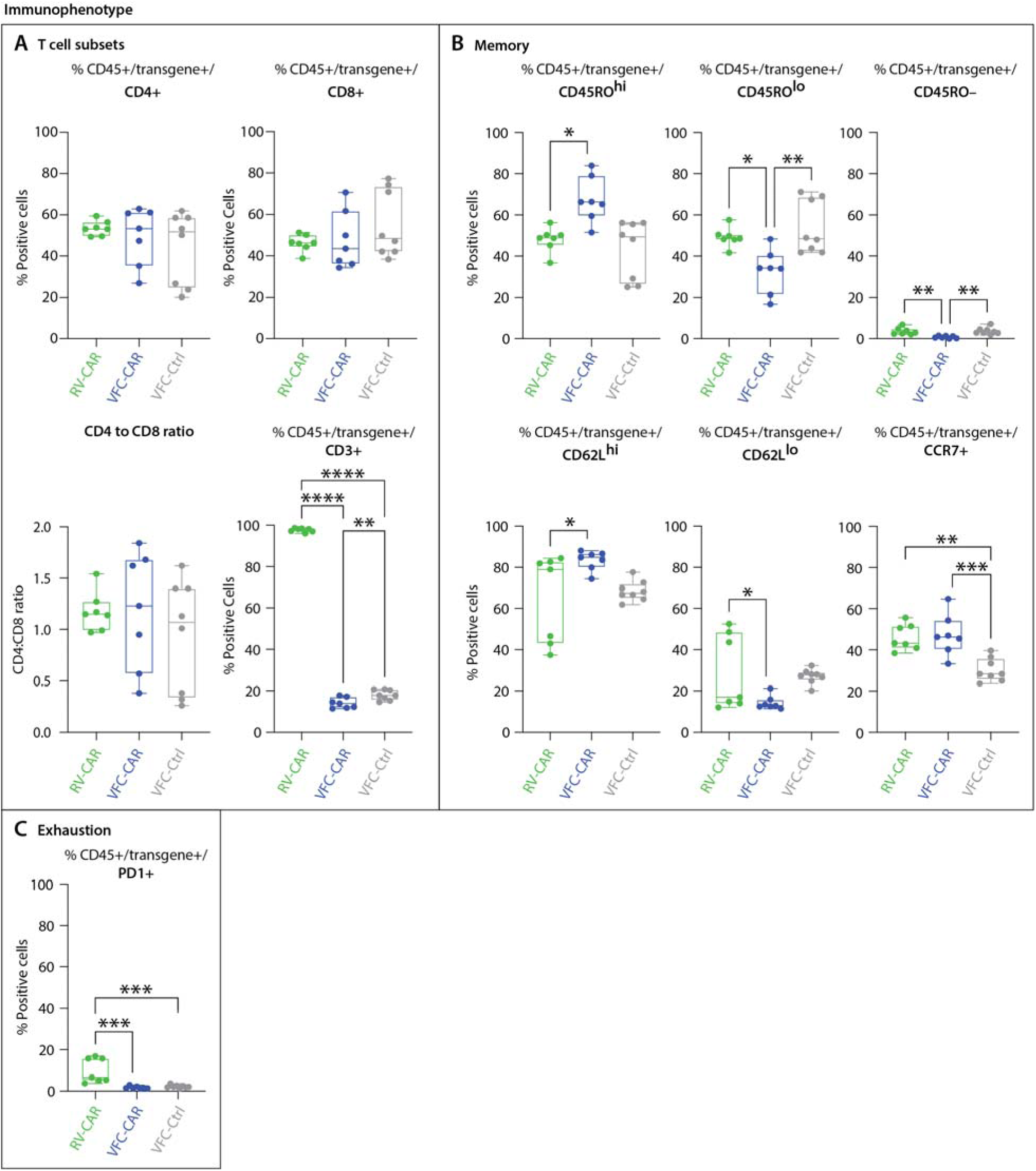
Immunophenotyping profile of VFC-CAR T products. Cells were assayed by spectral cytometry with a 21-color immunophenotyping panel on day 10 of manufacturing. (A) No significant differences were observed in CD4 and CD8 expression, or the CD4:CD8 ratio across cell types. CD3 was significantly decreased in *TRAC* edited products. (B) VFC-CAR T cells showed elevated levels of CD45RO^hi^ cells relative to RV-CAR T cells, and elevated levels of CD62L^hi^ cells; no significant difference was observed for CCR7, a third marker of central memory. (C) RV-CAR T cells showed elevated levels of the exhaustion marker PD-1 relative to VFC T cells. For all panels, cells were gated on CD45+ transgene+ cells (either CAR or mCherry). Gating strategies are shown in online supplemental figures S3. Additional markers are shown in online supplemental **figure S4**. VFC-CAR (blue) N=7; RV-CAR (green) N=7; VFC-Ctrl (gray) N=8 across two donors. Significance was determined by ordinary one-way ANOVA; * indicates p≤0.05; ** indicates p≤0.01; *** indicates p≤0.001; **** indicates p≤0.0001.

We also probed the activation-associated marker HLA-DR and five exhaustion-associated markers: PD1, LAG3, TIM3, TIGIT, and CD39. High levels of HLA-DR expression were seen in all groups, demonstrating proper stimulation from the IL-2 and activator present in the cell culture media. Of the exhaustion markers, only PD1 showed a significant difference across sample types, with elevated expression in RV-CAR T cells relative to either VFC-CAR or VFC-Ctrl products (**figure 4C**). While not fully exhausted at the end of the culture process, as evidenced by the ability to secrete more cytokine after antigen stimulation (**figure 3A**), RV-CAR T cells have progressed closer to terminal differentiation relative to VFC-CAR T cells. Other markers profiled included CD4, CD8, CD27, CD28, and CXCR3, and showed minimal or no differences among cell products. All products had negligible expression of the senescence marker CD57 (**online supplemental figure S4A**).

### Single cell memory- and exhaustion-associated transcriptional signatures of VFC-CAR products

To further characterize the phenotypic differences between RV-CAR, VFC-CAR, and VFC-Ctrl T cells, we performed single-cell RNA-sequencing (scRNA-seq) on 79,317 cells (post-quality control) from two different donors, both at the end of the manufacturing process and after 24 hours of co-culture with GD2+ CHLA20 neuroblastoma cells (**figure 5A**). We observed no significant donor-specific or batch effects, as indicated by gross clustering patterns in the combined data set (**online supplemental figure S5A**). Further, no significant changes in transcript levels were found for genes at or within 5 kb of off-target sites predicted by CHANGE-seq, indicating that any potential genomic disruptions at these sites did not lead to immediately detectable changes in proximal transcripts. To distinguish edited transgene-positive and transgene-negative cells within each sample, we aligned reads to custom reference genomes containing an added sequence mapping to the CAR or mCherry transgenes. Subsequent transcriptional analyses were carried out on transgene-positive cells only within each sample (21,068 total transgene+ cells). Untransfected cells were also profiled for one donor. Only 5/9623 untransfected cells contained any reads that mapped to the CAR transgene, indicating a false-positive rate of 0.05% in identifying transgene-positive cells.

**Figure 5.**
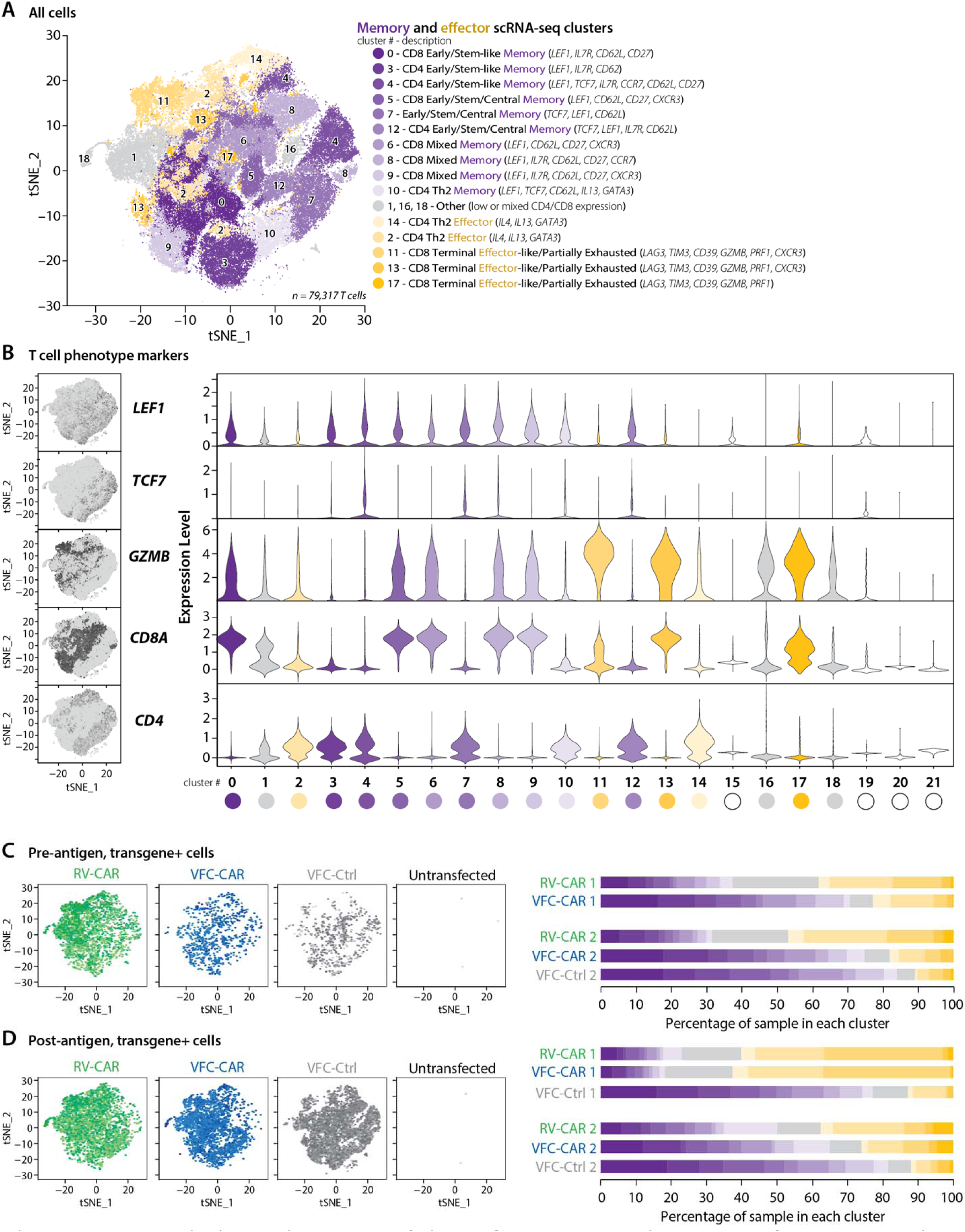
Transcriptional signatures of single CAR T cells prior to and after target antigen exposure. (A) tSNE projection of single cell RNA-seq data from 15 samples of manufactured cell products, both pre- and post-antigen exposure; 79,317 single cells from RV-CAR, VFC-CAR, VFC-Ctrl and untransfected T cell products are shown. (B) Feature plots showing distribution of *CD4*, *CD8A*, the stemness-associated markers *TCF7* and *LEF1*, and the effector-associated marker *GZMB*. At right are the expression levels of single cells within each of the clusters for these markers. (C, D) Proportion of transgene+ cells from all pre-antigen samples (C) and post-antigen samples (D) within each annotated cluster. Each color represents a different cluster, shown in A; purple clusters are memory-associated; yellow clusters are effector-associated; grey clusters could not be identified as pure T cell clusters due to a mix or lack of robust CD4/CD8 expression. At left, the distribution of transgene+ cells within the tSNE space in A are shown. 5/9623 untransfected cells featured reads mapping to the CAR or mCherry transgenes, indicating a false-positive rate of identifying transgene positive cells at 0.05%.

We performed tSNE dimensionality reduction and graph-based clustering on all cells and identified clusters with gene expression patterns associated with various phenotypes, including memory-like and effector-like populations across CD4 and CD8 subsets (**figure 5, online supplemental figure S6**). Using established unbiased clustering methods^43^, we identified 22 total clusters, of which 18 expressed T cell markers. The remaining 4 cancer-associated clusters arise residual cancer cells from our post-antigen samples and were removed from downstream analysis. The 18 T cell clusters exhibited more gradations of more stem-like, central and effector-like memory T cell formation, as well as populations with a mix of these phenotypes. Cluster identification was accomplished by assessing relative gene expression of markers associated with various T cell subtypes (**online supplemental figures S7-S9**) based on prior studies of human T cells^40,44,45^, including CAR T cells^46^. RNA-seq expression patterns for the protein markers profiled via immunophenotyping (**figure 4, S4**) are shown in **online supplemental figure S10**, and generally show concordance with protein-level expression. Consistent with our immunophenotyping results, we identified cells in transitional states of memory formation, expressing various combinations of markers associated with stemness, central memory, and effector function^40,44–47^ such as TCF7, LEF1 (**figure 5B**), CD95, IL7R, CD62L, CCR7, and others (**online supplemental figure S7**). Notably, some CD4 T cells were classified as belonging to memory or effector Th2 subsets (**online supplemental figures S7, S8A**). We also distinguished cells with effector-like phenotypes (e.g. high Granzyme B, *GZMB*; **figure 5B** **and online supplemental figure S8B**). These cells had some expression of exhaustion markers including LAG3, TIM3, and CD39, although PD1 transcripts were notably absent (**online supplemental figure S9**). All products contained a heterogeneous population of cells that were progressing toward but had not yet reached terminal differentiation and exhaustion.

The distribution of individual transgene+ cells within each cluster varied across the T cell samples (**figure 5C, D**). Prior to GD2 antigen exposure, 72% of VFC-CAR and 84% of VFC-Ctrl T cells were in clusters with a memory-like phenotype (Early/stem-like/central/mixed memory, clusters 0, 3, 4, 5, 6, 7, 8, 9, 10, 12 in **figure 5A**), while only 34% of RV-CAR T cells did so. 42% of RV-CAR T cells at harvest had effector-like or partially exhausted phenotypes (clusters 2, 11, 13, 14, 17 in **figure 5A**), while 21% of VFC-CAR T cells and 18% of VFC-Ctrl T cells fell into these clusters (**figure 5C**). As expected, the skew towards a memory-like phenotype did not persist for VFC-CAR or RV-CAR T cells after 24 hours of co-culture with GD2+ neuroblastoma cells *in vitro*. The distribution of transgene+ cells within each cluster shifted such that 44% and 48% of VFC-CAR and RV-CAR T cells, respectively, expressed effector-like transcriptional signatures after coculture, while 41% of VFC-CAR and 36% of RV-CAR T cells retained memory-associated transcriptional signatures. In contrast, only 12% of VFC-Ctrl cells expressed effector-like transcriptional signatures after coculture, while 79% of VFC-Ctrl cells retained a memory-associated transcriptional signatures (**figure 5D**). Based on this single cell analysis, individual VFC-CAR and RV-CAR T cells can mount a robust effector response, while individual VFC-Ctrl cells lacking either a CAR or TCR retain their less-differentiated phenotype when exposed to cancer cells.

### VFC-CAR T cells demonstrate potent *in vitro* killing of GD2-positive cancer cells

After characterizing cellular phenotypes and gene expression at the end of the manufacturing process, we measured the *in vitro* potency of VFC-CAR T cells against two GD2-positive solid tumors: CHLA20 neuroblastoma and M21 melanoma (**figure 6A**). We performed a fluorescence-based cytotoxicity assay measuring loss of expression from fluorescently labeled cancer cells over time (**figure 6B**), and IncuCyte live cell analysis at 2-hour intervals over a 48 hour period (**figure 6C**). We observed potent killing at a 5:1 effector:target ratio for both VFC-CAR and RV-CAR T cells, for both assays. These results corroborate our finding that VFC-CAR T cells produce proinflammatory cytokines and upregulate a cytotoxicity-associated gene signature at levels comparable to RV-CAR T cells and demonstrate potent target cell killing for multiple GD2+ cancers of variable origin.

**Figure 6.**
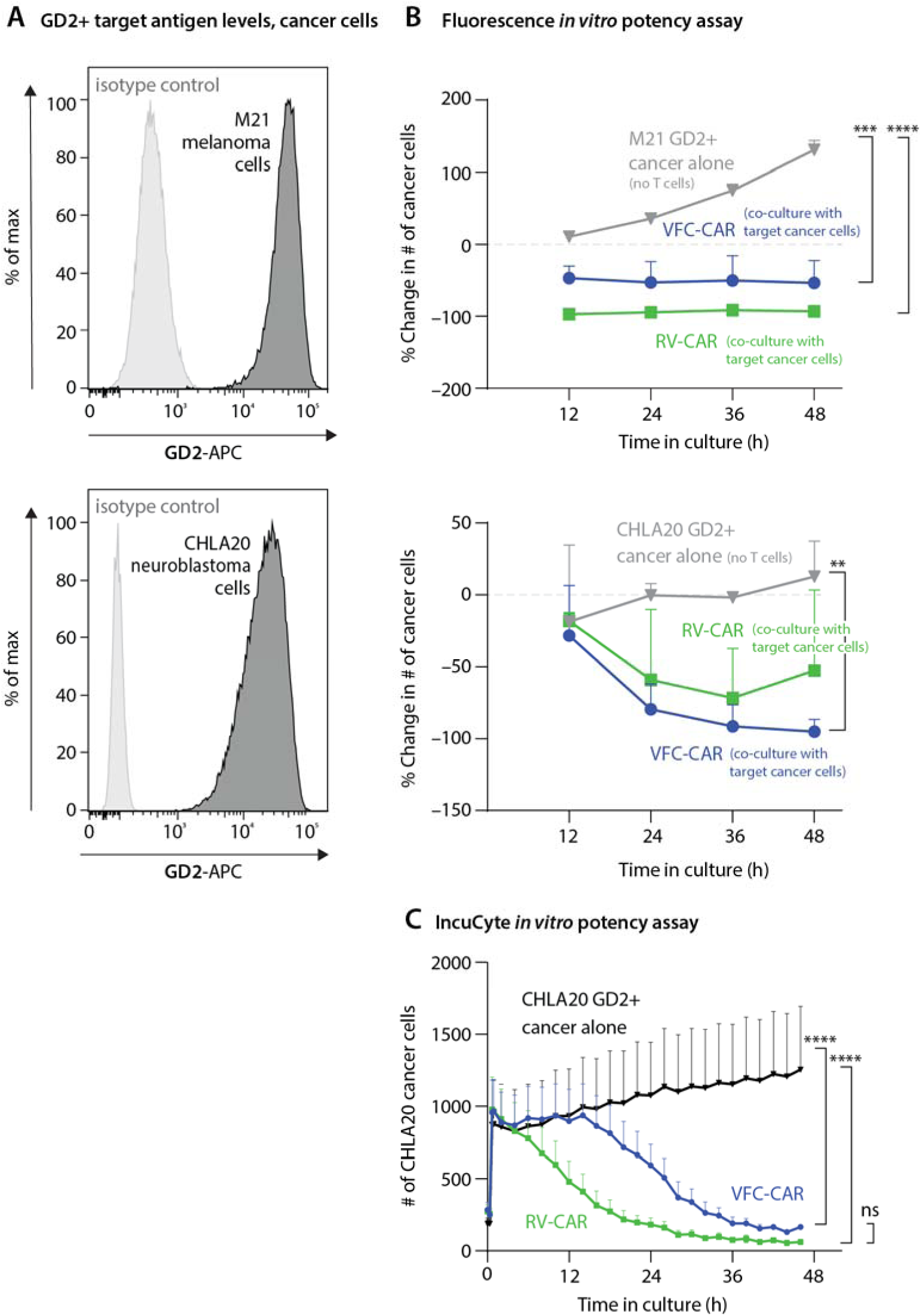
VFC-CAR T cells demonstrate robust *in vitro* killing of GD2+ cancer cells. (A) Flow cytometry histograms show GD2 surface expression on M21 and CHLA20 cell lines (black) compared to isotype controls (gray). (B) Graphs show the cytotoxic action of VFC-CAR and RV-CAR T cells against two GD2-positive tumor cell lines, CHLA20 and M21, containing a stably integrated H2B-mCherry fluorescent transgene. Cytotoxicity was measured as the change in the number of mCherry-positive objects for each image. The assay was performed using cells manufactured from one donor. (C) IncuCyte *in vitro* assay of T cell potency, averaged across four donors. AnnexinV was added as a marker of cell death; y-axis shows Akaluc-GFP-positive cancer cells in each well of a 96-well plate. The ratio of T cells to cancer cells is 5:1. The consistent decrease in CHLA20 cells after 15 hours indicates high potency of both VFC-CAR and RV-CAR T cells. VFC-CAR (blue) N=12; RV-CAR (green) N=12; CHLA20 neuroblastoma alone (black) N=9. *indicates p≤0.05; ** indicates p≤0.01; *** indicates p≤0.001; **** indicates p≤0.0001.

### VFC-CAR T cells induce regression of GD2-positive neuroblastoma *in vivo*

Because important clinical cell behaviors like homing, persistence and cytotoxicity within a tumor microenvironment cannot be easily assessed *in vitro*, we rigorously assessed CAR T cell potency *in vivo* in an established human GD2+ neuroblastoma xenograft model. After 9 total days of culture, multiple replicate wells of RV-CAR, VFC-CAR, or VFC-Ctrl T cells were pooled for injection into NOD-SCID-γc^-/-^ (NSG) mice (Jackson Laboratory). Ten million T cells were delivered via tail vein injection to each NSG mouse with an established luciferase-expressing CHLA20 neuroblastoma solid tumor identified by bioluminescence (**figure 7A**). Tumor sizes were quantified over time by IVIS imaging and digital caliper. Both CAR-treated cohorts showed robust tumor regression in the first 3 weeks post-infusion (**figure 7B, online supplemental figure S11A, B**). These cohorts also showed significantly improved survival relative to VFC-Ctrl-treated mice; however, there was no significant difference in survival between VFC-CAR and RV-CAR treated mice by day 80 (p-value=0.4099, n.s.; **figure 7C**). The percentage of CAR-positive cells per dose was lower in VFC-CAR T cells versus RV-CAR T cells (18% vs. 40%), which may have contributed to a slight decrease in complete remission rates (5/8 RV-CAR vs. 4/9 VFC-CAR). Inconsistencies in initial tumor burden may have affected remission. None of the VFC-Ctrl mice showed tumor regression, and all seven mice died of tumor progression by day 60, implying that disease control was antigen-specific. We also assessed persistence, memory and exhaustion phenotypes in CAR T cells isolated from the spleens and tumors of CHLA20-bearing mice as they reached euthanasia criteria. RV and VFC-CAR T cells persisted in both the spleens and tumors of the treated mice, but not for VFC-Ctrl T cell treatments. For the non-responding mice, VFC-CAR and RV-CAR T cells were robustly detected in tumors upon euthanasia of the treated mice, but almost no VFC-Ctrl T cells could be detected in their tumor counterparts (12%, 29%, and 0.058±0.046% human CD45+ cells within the tumor for VFC-CAR, RV-CAR, and VFC-Ctrl, respectively, **figure 7E**). These data indicate successful trafficking of VFC-CAR T cells to the tumor. (**figure 7D, E, online supplemental figure S11C, D**).

**Figure 7.**
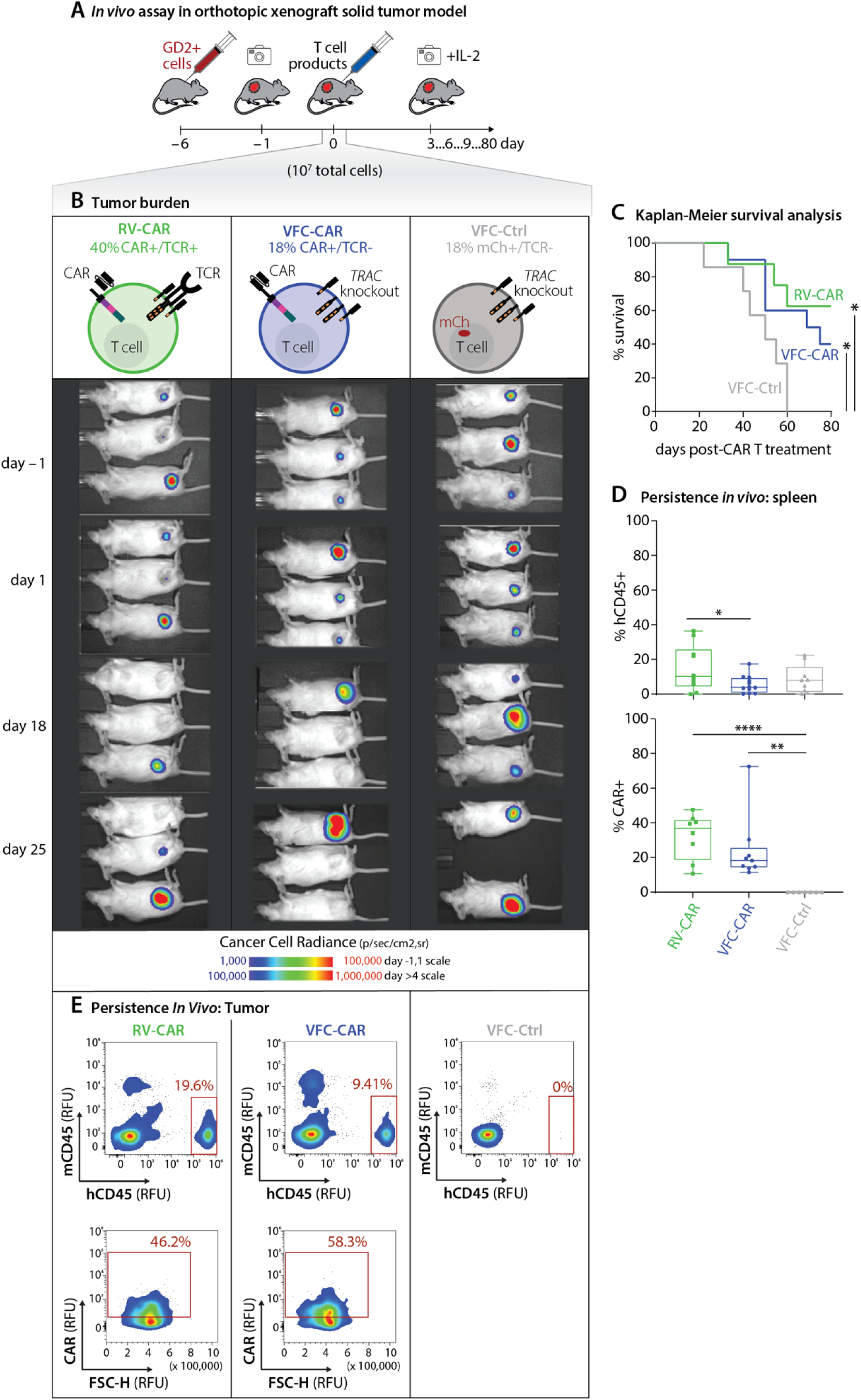
VFC-CAR T cells induce robust regression of GD2+ neuroblastoma solid tumors *in vivo* with high persistence. (A) Schematic of the *in vivo* mouse dosing strategy using NSG mice harboring GD2-positive CHLA20 neuroblastoma tumors. (B) Representative IVIS images of NSG mice with CHLA20 tumors that were treated with either 10 million VFC-CAR, RV-CAR, or VFC-Ctrl T cells. (C) Kaplan-Meyer survival curve for mice. VFC-CAR (blue) N=10; RV-CAR (green) N=8; VFC-Ctrl (gray) N=7. (D) Box plots show presence of CAR+CD45+ human T cells in mouse spleens, as measured by flow cytometry. (E) Flow cytometry plots show that human CD45+CAR+ VFC-CAR and RV-CAR T cells are found in tumors, but VFC-Ctrl cells are not.

Finally, to test the hypothesis that variations in gene transfer efficiency affected potency *in vivo*, we matched the absolute number of CAR-positive cells infused between virus-free and retroviral products in a separate xenograft study (**figure 8A**). NSG mice harboring human GD2+ CHLA20 xenograft tumors were infused with 10 million cells from three different T cell products. The percentage of CAR positive cells were equivalent (40%) in both the VFC-CAR and RV-CAR products. The VFC-Ctrl product had 38% transgene-positive cells. After one month, all four mice treated with RV-CART cells had higher adverse clinical scores indicative of xenogenic graft-vs-host-disease (xeno-GvHD; **figure 8A, D**). The lack of xeno-GvHD in the mice treated with VFC-CAR and VFC-Ctrl products indicates a functional knockout of TCR signaling by our CRISPR-Cas9 editing strategy. In contrast, three of the four mice treated with VFC-CAR products were event-free (no palpable tumor or GvHD) and survived past 96 days (**figure 8B-D**). We again assessed persistence, memory and exhaustion phenotypes in human lymphocytes recovered from the spleen and tumors of CHLA20-bearing mice as mice reached euthanasia criteria, up to 100 days after the initial T cell infusion. CAR+ or control mCherry+ T cells persisted in the spleens for all products (6.7±11.6%, 40±28%, and 26±12.4% human CD45+ cells within the spleen for VFC-CAR, RV-CAR and VFC-Ctrl, respectively). Of these cells, RV-CAR cells expressed higher levels of the exhaustion markers PD-1, LAG-3, and/or TIM-3 relative to VFC-CAR and VFC-Ctrl cells (**figure 8E**). Significantly higher numbers of RV-CAR T cells were differentiated toward effector memory and terminal effector cell states *in vivo* (**figure 8F**). We also observed elevated levels of the memory-associated proteins CCR7 and CD62L, and significantly lower expression of CD95 in VFC-CAR T cells relative to RV-CAR T cells (**online supplemental figure S11D**). These results *in vivo* mirror the significant skew toward effector phenotypes in RV-CAR cells seen *in vitro* with single cell RNA-seq and immunophenotyping assays. Altogether, these findings demonstrate comparable potency of VFC-CAR T cells to standard RV-CAR T cells, establishing the potential clinical relevance of VFC-CAR T cells for treating solid tumors.

**Figure 8.**
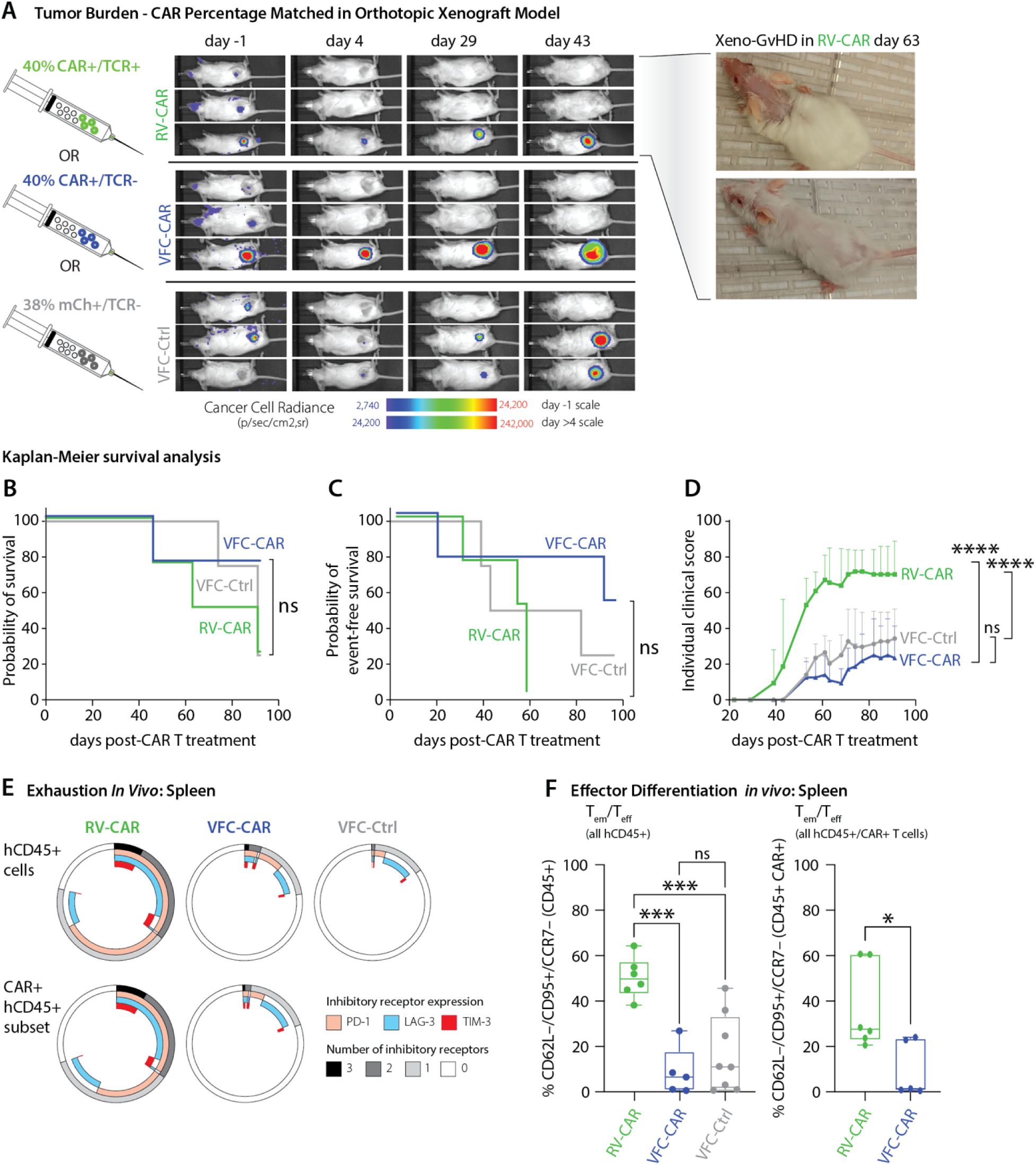
Virus-free CART cells exhibit *in vivo* potency against GD2+ solid tumors with high event-free survival and low exhaustion. (A, left) Representative IVIS images of NSG mice with CHLA20 tumors that were treated with either 10 million VFC-CAR, RV-CAR, or VFC-Ctrl T cells. VFC-CAR and RV-CAR products were 40% CAR-positive for a total dose of 4 million CAR+ cells per mouse. VFC-Ctrl products were 38% mCherry-positive for a total dose of 3.8 million transgene+ cells per mouse. GD2+ solid tumors were established in the side flank of each mouse as detected by IVIS imaging at day -1. At day 0, three different CART products as shown below were infused into the tail vein. (A, right) Pictures of RV-CAR T-treated mice showing xeno-GvHD symptoms from the intact TCR function within the RV-CAR T cells. None of the mice infused with VFC products displayed signs of xeno-GvHD. (B) Kaplan-Meier curve for total probability of survival. VFC-CAR (blue) N=4; RV-CAR (green) N=4; VFC-Ctrl (gray) N=4. (C) Kaplan-Meier curve for probability of event-free survival, defined as the absence of a palpable tumor or development of an individual clinical score of 4 or above. (D) Individual adverse clinical score of each mouse treated. Higher score indicates more adverse symptoms observed in the mice, such as elevated weight loss, hunched posture, ruffled fur, scaly or flaky skin, and decreased activity. (E) Donut plots show expression of exhaustion markers associated detected within T cells collected from mouse spleens. RV-CAR, N= 6; VFC-CAR, N=7, VFC-Ctrl, N=6. (F) T cell differentiation immunophenotypes detected within mouse spleens. RV-CAR T cells showed significantly higher proportions of more differentiated effector memory (T_em_) and terminal effector (T_eff_) T cells relative to VFC T cells.

## Discussion

Historically, CAR T cells have exhibited frustratingly limited success against solid tumors. While anti-GD2 CAR T cells were the first to mediate regression of a solid tumor^48^ in patients, the effects ultimately were not durable due in part to poor T cell persistence^27^. The third generation CAR used in this study failed to mediate meaningful anti-tumor responses in patients when delivered as a retroviral construct^49^; therefore, we sought to determine whether a TRAC-CAR replacement strategy, previously shown to be successful in the context of a CD19 CAR, could improve outcomes. Extensive work has focused on overcoming the immunosuppressive tumor microenvironment; however, there is an urgent need for new engineering strategies to make the cell product itself more potent, whether through armored CARs, T cell selection procedures, combinatorial therapies, or other approaches^50–52^. Leveraging prior work on hematological malignancies where anti-CD19 AAV-CAR T cells were generated using AAV and Cas9^17^, we develop a completely virus-free workflow that can accommodate a large CAR template (∼3.4kb) targeting a solid tumor antigen, GD2. Our findings suggest that manufacturing high-quality and defined genome-edited, VFC-CAR T cells to treat solid tumors is feasible.

Our manufacturing process produced similar yields across five donors and resulted in stable, genomically-integrated, durable CAR expression (>100 days *in vivo*) without the use of any viral vectors or animal-derived components during gene transfer or scale up. The decrease in T cell viability linked to electroporation^53^ is transient in our workflow with cells recovering to >80% viability just one week after electroporation, satisfying typical regulatory specifications^54^. Our use of high-density culture to improve T cell aggregation may stimulate pro-survival cell-cell signaling to overcome stress arising from electroporation. Furthermore, cell proliferation is likely to have an added benefit, as homologous recombination is active in the S and G2 phases of the cell cycle and increases in HDR have been observed in cycling cells^55^. We also modify the manufacturing process to generate the HDR repair template by performing two sequential Solid Phase Reversible Immobilization-based purification steps on the PCR products; this procedure concentrates the template. We do not rely on excipients to increase editing efficiency, which have been proposed recently^56^, and thereby provide a streamlined gene transfer process.

In retroviral and transposon-based CAR T products, vector copy numbers can vary,^57,58^ and genomic integration is scattered across >10,000 sites in the human genome^3^. CAR expression in retroviral and transposon-based products therefore can be affected by both the copy number and various chromatin contexts of each vector integrant across the various cells in a product. In contrast, our strategy inserts the CAR at a single site (*TRAC*) at a copy number of 1 or 2, where the CAR transgene is driven by the endogenous *TRAC* promoter. CRISPR strategies with AAV donor templates can integrate, at 5-20% frequencies, the entire AAV vector into the Cas9-induced DNA double-strand break^59^ and therefore may disrupt *TRAC* regulation. Hence, variable CAR expression within a cell product may depend on the degree of AAV vector integration. Usage of a virus-free CRISPR strategy can reduce variability in CAR expression (**figure 1F**). Undesired genomic alterations and adverse events arising from genome editing are low: the on-target specificity of our editor is above average when compared to many other editing strategies^35^, with no detectable transcriptional changes in genes at or proximal to predicted off-target sites in the edited cells. Our strategy to use Cas9 as RNPs versus encoded on mRNAs^17,60–62^ reduces the lifetime of active Cas9 proteins within the cell (hours for RNP versus days for mRNA). Transient induction of DNA double-strand breaks by Cas9 RNPs in our strategy likely contributes to the high specificity of genomic modifications to the *TRAC* on-target site, as increased off-target effects have been previously seen with prolonged presence of Cas9 within cells^63^. Our virus-free editing strategy can accommodate third-generation CAR sequences requiring the use of 3-4 kb nucleic acid templates. Transgene knockin with templates greater than 2 kb has historically been inefficient, although a recent report demonstrated efficient knockin of 2-3.6 kb templates^31^.

There is a paucity of knowledge regarding the signaling effects of CAR expression in T cell products^64^. We show evidence of decreased receptor signaling in the VFC-CAR T cell product at the level of the secretome and CD3ζ phosphorylation. Our results indicate that, in addition to altering the design of the CAR itself, the locus of insertion and the absence of TCR expression can affect receptor signaling. Reduced CAR and TCR tonic signaling during manufacturing could be notably important for allogeneic workflows involving stem cell sources (e.g., induced pluripotent stem cells^65–67^, hematopoietic stem cells^68^, umbilical cord blood^69^, etc.), where developmental signaling for proper specification towards effector cell types may be disrupted by receptor signaling during differentiation. We show evidence of heterogeneity in differentiation state at the protein and transcriptomic levels, which may in part be influenced by changes in CAR and TCR signaling throughout manufacture. Our large-scale scRNA-seq dataset profiling CAR T cells with varied receptor signaling profiles, both with and without antigen exposure and across multiple donors in this study, could be a useful resource for analyzing the effects of CAR transgenes within human immune cell products. This scRNA-seq dataset is the first such resource profiling CRISPR-generated CAR T cell products, to our knowledge.

The phenotype of the VFC-CAR T cell product could be advantageous for future clinical purposes. Cytokine production is lower in VFC-CAR T cells prior to antigen exposure, but equivalent to or higher than RV-CAR T cells post-antigen exposure, suggesting a higher dynamic range of antigen-driven potency. The VFC-CAR T cells also demonstrate increased expression of various memory-associated proteins, including CD62L and CD45RO, and decreased expression of the exhaustion marker PD1, relative to RV-CAR T cells. Prior to cognate antigen exposure, *TRAC* knockout may play a role in memory formation and maintenance of a less differentiated phenotype, as measured at the transcriptional level. This quality attribute is directly correlated with improved rates of durable remission with CD19 CAR T cells for hematologic malignancies^19,70^. Prior work with hematopoietic stem/progenitor cells indicated that the AAV template itself elicited both immune and stress responses, along with transcriptional downregulation of cell cycle processes that could interfere with stem cell maintenance^71^. Switchable anti-CD19 CARs have demonstrated increased CD62L+ memory formation upon turning off CAR signaling, indicating that prolonged or tonic CAR signaling can interfere with memory formation^72^. These studies are consistent with prior work where overstimulation of TCR signaling and CD28 co-stimulatory signaling can affect unmodified T cell differentiation *in vivo*, as memory responses *in vivo* with unmodified T cells are formed through acute, high-load antigen stimulation followed by a “rest” phase^73^. Future studies may also reveal additional mechanistic connections between these observations.

After injection into a GD2-positive human neuroblastoma xenograft model, VFC-CAR T cells induce strong regression of solid tumors compared to mock-edited T cells, and at levels comparable to RV-CAR T cells. In our first *in vivo* experiment, comparable tumor regression rates occurred despite the substantially lower proportion of CAR+ cells in the VFC-CAR product. In our second *in vivo* experiment, we observed that cell-for-cell, an VFC-CAR T cell product could lead to a potent *in vivo* response with longer event-free survival. The TCR knockout by our editing strategy is functionally validated in this second study, as xenogenic GvHD was significantly delayed or eliminated altogether for the mice treated VFC-CAR and VFC-Ctrl products.

Finally, relative to conventional T cell manufacturing, a virus-free manufacturing process could have several advantages at clinical scale. First, it could reduce batch-to-batch variability, supply chain challenges, and costs associated with vector production^8,74^. Second, it could alleviate a number of regulatory considerations related to the need for monitoring replication competency of the vector and the levels of xenogeneic components in the clinical cell product, notably plasmid DNA and serum during the gene transfer and scale up process that can introduce infectious agents or toxic components^75^. Third, it could eliminate the potential for integration of viral elements into the human genome, which can generate a high degree of gene perturbation, up to 10^4^-10^5^ different insertional sites within a single product^73^. Integration of the vector, in particular, presents risks of insertional oncogenesis^76^, transgene silencing or overexpression, and adverse immune response to the vector, which could result in the rejection of therapeutic cells. While off-target analysis of genome editors is necessary for any clinical translation of our approach, there are now many experimental and computational tools that can readily be used for this purpose^35,77^ and next-generation high-fidelity Cas9 enzymes^78^ could be used to further decrease the potential for any off-target effects. Overall, a virus-free genome editing workflow has high potential to enable the rapid and flexible manufacture of highly defined and highly potent CAR T cell products for the treatment of solid tumors.

## Methods

### Cell lines

CHLA20 human neuroblastoma cells were a gift from Dr. Mario Otto and M21 human melanoma cells were a gift from Dr. Paul Sondel (University of Wisconsin-Madison). These cells were maintained in Dulbecco’s Modified Eagle Medium high glucose (Gibco) supplemented with 10% Fetal Bovine Serum (Gibco) and 1% Penicillin-Streptomycin (Gibco). H2B-mCherry-positive lines of M21 and CHLA20 cells were generated via lipofection for the fluorescence *in vitro* assay. AkaLUC-GFP CHLA20 cells were a gift from Dr. James Thomson (Morgridge Institute for Research). Phoenix cells (ATCC) for viral preparation were maintained in DMEM (high glucose) supplemented with 10% Fetal Bovine Serum (Gibco) and selected using 1 µg/mL diphtheria toxin (Cayman Biologics) and 300 µg/mL hygromycin (Thermo Fisher Scientific) prior to use. Selection for transgene positive cells was confirmed by flow cytometry for mouse Lyt2 expression (Biolegend) (>70%+). 3T3 cells were maintained in Dulbecco’s Modified Eagle Medium (Gibco) supplemented with 10% Fetal Bovine Serum (Gibco) and 1% Penicillin-Streptomycin (Gibco). Cell authentication was performed using short tandem repeat analysis (Idexx BioAnalytics, Westbrook, ME) and per ATCC guidelines using morphology, growth curves, and *Mycoplasma* testing within 6 months of use with the e-Myco mycoplasma PCR detection kit (iNtRON Biotechnology Inc, Boca Raton, FL). Cell lines were maintained in culture at 37°C in 5% CO_2_.

### Plasmid constructs

VFC-CAR: A 2kb region surrounding the *TRAC* locus was amplified by PCR from human genomic DNA and cloned into a pCR blunt II TOPO backbone (Thermo Fisher Scientific). The CAR transgene from a pSFG.iCasp9.2A.14G2A-CD28-OX40-CD3ζ RV-CAR plasmid (gift from Dr. Malcolm Brenner, Baylor College of Medicine) was then cloned into the TOPO *TRAC* vector using Gibson Assembly (New England Biolabs (NEB)). The plasmid sequence was verified by Sanger sequencing. The VFC-Ctrl (mCherry) construct was designed in house, synthesized, and sequence-verified (GenScript). All plasmids were grown in 5-alpha competent *E. coli* (NEB) and purified using the PureYield MidiPrep system (Promega).

### Double-stranded DNA HDR template production

Plasmid donors were used as PCR templates for VFC products. In brief, VFC-CAR and VFC-Ctrl plasmids were MidiPrepped using the PureYield MidiPrep system (Promega). PCR amplicons were generated from plasmid templates using Q5 Hot Start Polymerase (NEB) and pooled into 100 µl reactions for Solid Phase Reversible Immobilization (SPRI) cleanup (1X) using AMPure XP beads according to the manufacturer’s instructions (Beckman Coulter). Each 100 µl starting product was eluted into 5 µl of water. Bead incubation and separation times were increased to 5 minutes, and elution time was increased to 15 minutes at 37°C to improve yield. PCR products from round 1 cleanup were pooled and subjected to a second round of SPRI cleanup (1X) to increase total concentration; round 2 elution volume was 20% of round 1 input volume. Template concentration and purity was quantified using NanoDrop 2000 and Qubit dsDNA BR Assays (Thermo Fisher Scientific), and templates were diluted in water to an exact concentration of 2 µg/µl according to Qubit measurements.

### SpCas9 RNP preparation

RNPs were produced by complexing a two-component gRNA to SpCas9. In brief, tracrRNA and crRNA were ordered from IDT, suspended in nuclease-free duplex buffer at 100 µM, and stored in single-use aliquots at -80°C. tracrRNA and crRNA were thawed, and 1 µl of each component was mixed 1:1 by volume and annealed by incubation at 37°C for 30 minutes to form a 50 µM gRNA solution in individual aliquots for each electroporation replicate. Recombinant sNLS-SpCas9-sNLS Cas9 (Aldevron, 10 mg/ml, total 0.8 µl) was added to the complexed gRNA at a 1:1 molar ratio and incubated for 15 minutes at 37°C to form an RNP. Individual aliquots of RNPs were incubated for at least 30 seconds at room temperature with HDR templates for each sample prior to electroporation.

### Isolation of primary T cells from healthy donors

This study was approved by the Institutional Review Board of the University of Wisconsin-Madison (#2018-0103), and informed consent was obtained from all donors. Peripheral blood was drawn from healthy donors into sterile syringes containing heparin, and transferred to sterile 50 mL conical tubes. Primary human T cells were isolated using negative selection per the manufacturer’s instructions (RosetteSep Human T Cell Enrichment Cocktail, STEMCELL Technologies). T cells were counted using a Countess II FL Automated Cell Counter (Thermo Fisher Scientific) with 0.4% Trypan Blue viability stain (Thermo Fisher Scientific). T cells were cultured at a density of 1 million cells/mL in ImmunoCult-XF T cell Expansion Medium (STEMCELL) supplemented with 200 U/mL IL-2 (Peprotech) and stimulated with ImmunoCult Human CD3/CD28/CD2 T cell Activator (STEMCELL) immediately after isolation, per the manufacturer’s instructions.

### T cell culture

Bulk T cells were cultured in ImmunoCult-XF T cell Expansion Medium at an approximate density of 1 million cells/mL. In brief, T cells were stimulated with ImmunoCult Human CD3/CD28/CD2 T cell Activator (STEMCELL) for 2 days prior to electroporation. On day 3, (24 hours post-electroporation), VFC-CAR and VFC-Ctrl T cells were transferred without centrifugation to 1 mL of fresh culture medium (with 500 U/mL IL-2, no activator) and allowed to expand. T cells were passaged, counted, and adjusted to 1 million/mL in fresh medium + IL-2 on days 5 and 7 after isolation. RV-CAR T cells were spinoculated with the RV-CAR construct on day 3 and passaged on day 5 along with the VFC-CAR and VFC-Ctrl T cells. Prior to electroporation or spinoculation, the medium was supplemented with 200 U/mL IL-2; post-gene editing, medium was supplemented with 500 U/mL IL-2 (Peprotech).

### T cell nucleofection

RNPs and HDR templates were electroporated 2 days after T cell isolation and stimulation. During crRNA and tracrRNA incubation, T cells were centrifuged for 3 minutes at 200g and counted using a Countess II FL Automated Cell Counter with 0.4% Trypan Blue viability stain (Thermo Fisher). 1 million cells per replicate were aliquoted into 1.5 mL tubes. During the RNP complexation step (see RNP production), T cell aliquots were centrifuged for 10 min at 90g. During the spin step, 2 µl of HDR template (total 4 µg) per condition were aliquoted to PCR tubes, followed by RNPs (2.8 µl per well; pipette should be set to a higher volume to ensure complete expulsion of viscous solution). Templates and RNPs were incubated at room temperature for at least 30 seconds. After cell centrifugation, supernatants were removed by pipette, and cells were resuspended in 20 µl P3 buffer (Lonza), then transferred to PCR tubes containing RNPs and HDR templates, bringing the total volume per sample to 24 µl. Each sample was transferred directly to a 16 well electroporation cuvette. Typically, no more than 8 reactions were completed at a time to minimize the amount of time T cells spent in P3 buffer. T cells were electroporated with a Lonza 4D Nucleofector with X Unit using pulse code EH115. Immediately after electroporation, 80 µl of pre-warmed recovery medium with 500 U/mL IL-2 and 25 µl/mL ImmunoCult CD3/CD28/CD2 activator was added to each well of the cuvette. Cuvettes were rested at 37°C in the cell culture incubator for 15 minutes. After 15 minutes, cells were moved to 200 µl total volume of media with IL-2 and activator (see above) in a round bottom 96 well plate.

### Retrovirus production

CAR retrovirus was manufactured using Phoenix cells (ATCC). In brief, pSFG.iCasp9.2A.14G2A-CD28-OX40-CD3ζ plasmid was MidiPrepped using the PureYield MidiPrep system (Promega). One day prior to transfection, selected Phoenix cells were plated on 0.01% Poly-L-Lysine coated 15 cm dishes (Sigma Aldrich) at a density of 76,000 cells/cm^2^, or ∼65% confluency. On transfection day, media was replaced 1 hour prior to transfection of 10 µg pSFG.iCasp9.2A.14G2A-CD28-OX40-CD3ζ plasmid/plate using iMFectin according to the manufacturer’s instructions (GenDEPOT). Media was replaced 18-24 hours later with 10 mL of 50 mM HEPES buffered DMEM + 10% FBS (Gibco). 48 hours later, media was collected, stored at 4°C, and replaced. A second aliquot of media was collected 24 hours later; media aliquots were pooled and centrifuged for 10 min at 2000g to pellet contaminating cells, and supernatants were transferred to a clean conical tube. 1/3 volume Retro-X concentrator (Takara) was added, and supernatants were refrigerated at 4°C for 12-18 hours, then concentrated according to the manufacturer’s instructions. Viruses were tested on 3T3 cells prior to use; yields from one 15 cm dish were used for 5 replicate wells of 160,000 T cells per transduction. Viruses were either used immediately for T cell spinoculation or stored at -80°C in single use aliquots.

### Retroviral transduction

T cells for RV infection were cultured similarly to VFC-CAR and VFC-Ctrl T cells, with two exceptions: 1) T cells were passaged and resuspended without ImmunoCult CD2/CD28/CD3 activator on day 2 post-isolation, and spinoculated on Day 3. RV-CAR T cells returned to the regular passaging schedule on day 5 post-isolation. Prior to spinoculation, non-tissue culture treated 24 well plates were coated with Retronectin according to the manufacturer’s instructions (Takara/Clontech). On day 3 post-isolation, T cells were centrifuged at 200 g for 3 minutes, counted, and resuspended to a concentration of 200,000 cells/mL, then stored in the incubator until plates were prepared. Virus was added to retronectin-coated plates in a volume of 400 µl virus in ImmunoCult-XF Medium and centrifuged at 2000g for 2 hours at 32°C. 160,000 T cells in 800 µl were added to each well and spinoculated at 2000g for 60 minutes at 32°C, brake off. T cells were then transferred to the incubator and left undisturbed for two days.

### Flow cytometry and fluorescence-activated cell sorting

CAR was detected using 1A7 anti-14G2a idiotype antibody (gift from Paul Sondel) conjugated to APC with the Lightning-Link APC Antibody Labeling kit (Novus Biologicals). T cells were stained in BD Brilliant Stain Buffer (BD Biosciences). Flow cytometry was performed on an Attune NxT Flow cytometer (Thermo Fisher Scientific) and an Aurora Spectral Cytometer (Cytek), and fluorescence-activated cell sorting was performed on a FACS Aria (BD). T cells were stained and analyzed on day 7 of manufacture for CAR and TCR expression, and day 10 of manufacture for the full Aurora immunophenotyping panel, using fresh cells. Downstream analyses of all spectral cytometry data were performed in FCS Express 7 Software. All flow cytometry antibodies are listed in **online supplemental table S2.**

### In-out PCR

Genomic DNA was extracted from 100,000 cells per condition using DNA QuickExtract (Lucigen), and incubated at 65°C for 15 min, 68°C for 15 min, and 98°C for 10 min. Genomic integration of the CAR was confirmed by in-out PCR using a forward primer upstream of the *TRAC* left homology arm, and a reverse primer binding within the CAR sequence. Primer sequences are listed in **online supplemental table S3**. PCR was performed according to the manufacturer’s instructions using Q5 Hot Start Polymerase (NEB) using the following program: 98°C (30 s), 35 cycles of 98°C (10 s), 62°C (20 s), 72°C (2 min), and a final extension at 72°C (2 min).

### Next Generation Sequencing of genomic DNA

Indel formation at the *TRAC* locus was measured using Next Generation Sequencing (Illumina). Genomic PCR was performed according to the manufacturer’s instructions using Q5 Hot Start polymerase (NEB); primers are listed in **online supplemental table S3**. Products were purified using SPRI cleanup with AMPure XP beads (Beckman Coulter), and sequencing indices were added with a second round of PCR using indexing primers (Illumina), followed by a second SPRI cleanup. Samples were pooled and sequenced on an Illumina MiniSeq according to the manufacturer’s instructions. Analysis was performed using CRISPR RGEN (rgenome.net).

### Genome-wide, off-target analysis

Genomic DNA from human primary CD4^+^/CD8^+^ T cells was isolated using the Gentra Puregene Kit (Qiagen) according to the manufacturer’s instructions. CHANGE-seq was performed as previously described^23^. Briefly, purified genomic DNA was tagmented with a custom Tn5-transposome to an average length of 400 bp, followed by gap repair with Kapa HiFi HotStart Uracil+ DNA Polymerase (KAPA Biosystems) and Taq DNA ligase (NEB). Gap-repaired tagmented DNA was treated with USER enzyme (NEB) and T4 polynucleotide kinase (NEB). Intramolecular circularization of the DNA was performed with T4 DNA ligase (NEB) and residual linear DNA was degraded by a cocktail of exonucleases containing Plasmid-Safe ATP-dependent DNase (Lucigen), Lambda exonuclease (NEB) and Exonuclease I (NEB). *In vitro* cleavage reactions were performed with 125 ng of exonuclease-treated circularized DNA, 90 nM of SpCas9 protein (NEB), NEB buffer 3.1 (NEB) and 270 nM of sgRNA, in a 50 μL volume. Cleaved products were A-tailed, ligated with a hairpin adaptor (NEB), treated with USER enzyme (NEB) and amplified by PCR with barcoded universal primers NEBNext Multiplex Oligos for Illumina (NEB), using Kapa HiFi Polymerase (KAPA Biosystems). Libraries were quantified by qPCR (KAPA Biosystems) and sequenced with 151 bp paired-end reads on an Illumina NextSeq instrument. CHANGE-seq data analyses were performed using open-source CHANGE-seq analysis software (https://github.com/tsailabSJ/changeseq).

### Cytokine Analysis

Cytokine analysis was performed using a V-PLEX Proinflammatory Panel 1 Human Kit (Meso Scale Discovery, Catalog No K15049D-2) according to the manufacturer’s protocol. The following cytokines were measured: IFNγ, IL-1β IL-2, IL-4, IL-6, IL-8, IL-10, IL-12p70, IL-13, and TNF-α. In brief, media was collected from the final day of cell culture before injection into mice and flash frozen and stored at -80°C. For co-culture samples, 250,000 T cells were co-cultured with 50,000 cancer cells in 250 µl ImmunoCult XF T cell expansion medium for 24 hours prior to media collection. On the day of the assay, media was thawed and 50 µl of media was used to perform all measurements in duplicate. Figures were produced using GraphPad PRISM 8. Data were normalized by calculating cytokine production per cell based on the total concentration of cells calculated at media collection.

### Immunoblotting

Equivalent number of T cells (1×10e6) were lysed in Laemmli Sample Buffer with β-mercaptoethanol (Bio-Rad, CA). Total cell lysate for each sample were resolved on 12% SDS-PAGE gels and transferred to polyvinylidene fluoride membranes (Millipore, Billerica, MA). The membranes were blocked in LI-COR blocking buffer (LI-COR, NE), Immunoblotting was performed by incubating the membranes with anti-human CD247 (Mouse, BD Biosciences), anti-human CD247 pTyr^1^^42^ (Mouse, BD Biosciences), and anti-human GAPDH (Rabbit, Cell Signaling Tech, MA), according to the manufacturer’s recommendations. The membranes were then washed with TBST and incubated with fluorescent secondary antibodies (LI-COR, NE) and the immunoreactive bands were visualized using the Odyssey ^®^ CLx imaging system (LI-COR, NE).

### In Vitro Cytotoxicity Assays

For figure 6C 10,000 AkaLUC-GFP CHLA20 cells were seeded in triplicate per condition in a 96 well flat bottom plate. 48 hours later, 50,000 T cells were added to each well. 1 µl (0.05 µg) of CF® 594 Annexin V antibody (Biotium) was added to the wells. The plate was centrifuged at 100g for 1 minute and then placed in The IncuCyte® S3 Live-Cell Analysis System (Sartorius, Catalog No 4647), stored at 37°C, 5% CO_2_. Images were taken every 2 hours for 48 hours. Green object count was used to calculate the number of cancer cells in each well. Red object count was used to calculate the number of objects staining positive for Annexin V, an early apoptosis marker. Fluorescent images were analyzed with IncuCyte Base Analysis Software. *For* ***figure 6B***: 10,000 H2B-mCherry CHLA20 cells or 10,000 H2B-mCherry M21 cells were seeded in triplicate per condition in a 96 well flat bottom plate. 24 hours later, 50,000 T cells were added to each well. The 96 well plate was placed in a live cell imaging chamber at 37°C and 5% CO_2_ and imaged on a Nikon Epifluorescent scope, with images taken every 12 hours for 48 hours. The change in protocol was made in March 2020 due to institutional COVID-19 biosafety precautions.

### Single cell RNA sequencing

24 hours prior to assay, 200,000 AkaLUC-CHLA20 cells were plated in 12 well plates and cultured overnight. One week after electroporation (day 9 post-isolation), T cells were counted and pooled into a single bank for characterization studies (scRNA-seq, IncuCyte cytotoxicity assay and *in vivo* experiments). Media was aspirated from cancer cells, and 1 million T cells in ImmunoCult-XF Medium + 500 U/mL IL-2 were seeded on the cancer cells, then cultured for 24 hours. A parallel T cell-only single culture (termed “pre-antigen”) was set up at the same density in a separate 12 well plate. The next day, co-cultured cells were trypsinized for donor 1 and washed off the plate with media, and cells were singularized with a 35 µM cell strainer prior to scRNA-seq (Corning). For donor 2, to improve the total purity of the T cell populations and remove contaminating cancer cells from analysis, co-culture cells were stained for CD45 and CAR, and FACS sorted into CD45^+^CAR^+^ and CD45^+^CAR^-^ fractions prior to sample submission. Cells were counted with a Countess II FL cell counter using trypan blue exclusion (Thermo Fisher Scientific), and samples were prepared for single cell RNA sequencing with the 10X Genomics 3’ kit (v3 chemistry) according to the manufacturer’s instructions. Libraries were sequenced using the Illumina NovaSeq 6000 system.

### Single cell RNA-sequencing analyses

*Alignment, Data Quality Control, Integration, Clustering, and Annotation:* FASTQ files were aligned with Cellranger v3.0.1 to custom reference genomes that included added sequences for the transgene(s) used in each culture condition (e.g., the *TRAC* VFC-CAR donor sequence, VFC-Ctrl mCherry donor sequence, etc.). Downstream analyses were performed using the Seurat package v4.0.1 in R software v4.0.3^43^. Several quality control measures were used to filter data prior to downstream processing. First, each dataset was filtered to include only cells with 200 or more unique genes, and genes expressed in three or more cells. To further preserve cell quality, any cells with greater than 15% mitochondrial RNA reads or less than 500 detected genes were also excluded. Additionally, maximum RNA and gene count thresholds were applied to each sample to filter out potential doublet or multiplet captures. Specific maximum thresholds were determined sample-by-sample, ranging from 80,000-100,000 and 7500-8500 for RNA counts and gene counts, respectively. Subsequent analyses were performed in Seurat using default settings, unless otherwise noted. Each sample was log-normalized (NormalizeData) and 2000 variable features were selected using FindVariableFeatures. All datasets were integrated using reference-based integration and Reciprocal Principal Component Analysis (RPCA) was used to identify integration anchors. In brief, the workflow was performed as follows: Each dataset was separately scaled (ScaleData) and dimensionally reduced using Principal Component Analysis (PCA) (RunPCA), setting the ‘features’ parameter in both functions equal to a vector containing all genes. Next, integration anchors were identified using the two untransfected controls as the references for anchor selection (FindIntegrationAnchors, reduction = ‘rpca’, dims = 1:50). Datasets were then integrated using all genes and the selected anchors (IntegrateData, dims = 1:50, features.integrate = all_genes variable). Following integration, the data were scaled (ScaleData) and dimensionally reduced with PCA (RunPCA) and T-distributed stochastic neighbor embedding (t-SNE) (RunTSNE; dims = 1:50). The data were then clustered (FindNeighbors, dims = 1:50; FindClusters). Cell-level annotations were derived using the Seurat multi-modal reference mapping pipeline with a human PBMC reference cell atlas^43^. One notable caveat of this pipeline is that all cells in the query dataset are forcibly mapped to the reference cell type that matches most closely. Consequently, it is conceivable that novel cell types present in the query dataset are lost to other cell labels. These cell-level annotations were then used to inform labeling of t-SNE clusters, in conjunction with manual review of canonical feature expression and differentially expressed genes for each cluster. Clusters 15, 19, 20, and 21 were largely composed of co-culture samples and lacked expression of canonical T cell markers. It was determined that these clusters represented contaminating CHLA20 cancer cells, which we subsequently removed from the dataset. Downstream comparisons of sample types were performed on transgene+ cells only. All analysis scripts will be deposited in the Saha Lab GitHub repository upon publication.

### *In vivo* human neuroblastoma xenograft mouse model

All animal experiments were approved by the University of Wisconsin-Madison Animal Care and Use Committee (ACUC). Male and female NSG mice (9-25 weeks old) were subcutaneously injected with 10 million AkaLUC-GFP CHLA20 human neuroblastoma cells in the side flank to establish tumors. Six days later (Day 0), established tumors were verified by bioluminescence with the PerkinElmer *In Vivo* Imaging System (IVIS), and 10 million T cells were injected through the tail vein into each mouse. Mice were followed for weight loss and overall survival. On imaging days, mice were sedated using isoflurane and received intraperitoneal injections of ∼120 mg/kg D-luciferin (GoldBio). Fifteen minutes later, mice were imaged via IVIS. Imaging was repeated every 3 to 4 days, starting 1 day before initial T cell injection (Day -1). Mice were injected with 100,000 IU of human IL-2 subcutaneously on day 0, day 4, and with each subsequent IVIS reading. In order to quantify the total flux in the IVIS images, a region of interest (ROI) was drawn around the bottom half of each mouse with the total flux being calculated by Living Image® software (PerkinElmer; Total flux = the radiance (photons/sec) in each pixel summed or integrated over the ROI area (cm^2^) x 4π. The absolute minimum total flux value was subtracted from each image to minimize background signal. For donors 1, 3, 4, and 5 mice were maintained until tumors reached 20mm in any dimension by digital caliper as defined by the ACUC.

### Flow cytometric analysis of splenic and tumor-infiltrating T cells

For donor 2, all mice were euthanized on day 25. Tumors and spleens were removed, mechanically dissociated, and passed through a Corning® 35µm cell strainer. Cell suspensions were centrifuged at 300g for 10 minutes, and then digested with ACK lysing buffer (Lonza). The cells were then washed and centrifuged at 300g for 10 minutes, and resuspended in 10 ml PBS, 10 µl of which was added to 10 ml of ISOTON® diluent and counted on the COULTER COUNTER® Z1 Series Particle Counter (Beckman Coulter). From this count, 1×10^6^ cells were added to flow cytometry tubes in staining buffer (PBS with 2% FBS) and stained with antibodies for hCD45, mCD45, scFV 14g2a CAR, and PD-1 (see **Supplementary Table 2 for antibody information**). The cells were then washed with PBS, centrifuged at 300g for 10 minutes, and 0.5ul of Ghost Dye™ Red 780 viability dye (Tonbo Biosciences) was added for 20 minutes at room temperature. Cells were then washed with staining buffer, spun down, and resuspended in 400 µl of staining buffer. Cells were run on an Attune™ NXT flow cytometer (Thermo Fisher Scientific). Subsequent analyses were performed using Flowjo™ software (BD). For donors 3 and 4, spleens and tumors were analyzed as mice reached euthanasia criteria.

### Statistical analysis

Unless otherwise specified, all analyses were performed using GraphPad Prism (v.8.0.1), and error bars represent mean ± SD; ns = p>=0.05, * for p<0.05, ** for p<0.01, *** for p<0.001,**** for p<0.0001. For Fig. 2b, error bars show SEM. Statistical analyses for cytokine data were performed using a two-tailed Mann-Whitney test in GraphPad Prism. Statistical analyses for flow cytometry data were performed using a one-way ANOVA test in GraphPad prism. All box plots show median (horizontal line), interquartile range (hinges), and smallest and largest values (whiskers). Statistical significance for **Fig. 7C** was calculated using the Mantel-Cox Test.

## Data Availability

The data that support the findings of this study will be made available in the public domain upon publication.

## Data Reporting

The Reporting Summary document includes information about the statistics, software, data, and sample preparation methods used for this study. For *in vivo* experiments, established tumor burden was verified by IVIS luciferase imaging prior to infusion. Mice were arranged according to tumor burden and distributed evenly across conditions. The experiments were not randomized and the investigators were not blinded during experiments and outcome assessment.

## Acknowledgments

We thank members of the Saha and Capitini labs and John Hosmer-Quint for helpful discussion and comments on the manuscript, the University of Wisconsin (UW) Carbone Cancer Center Flow Cytometry Laboratory for assistance with flow cytometry experiments, and Aldevron for technical support with Cas9 proteins. We thank Tyler Duellman and the University of Wisconsin-Madison Biotechnology Center Gene Expression Center the DNA Sequencing Facility for providing single nuclei library preparation and next generation sequencing services. We thank Malcolm Brenner (Baylor College of Medicine) for sharing the RV-CAR plasmid, Paul Sondel and the National Cancer Institute for 1A7 anti-14G2a antibody for detection of CAR expression, James Thomson and Jue Zhang (Morgridge Institute for Research) for the AkaLUC-GFP CHLA20 cancer line used for *in vivo* studies, and Sushmita Roy, Sunnie Grace McCalla, and Kurt Mueller for helpful discussions and assistance with scRNA-seq analysis. We thank Dan Cappabianca and Lauren Sarko for assistance with template production. We thank Adam Steinberg and Art for Science for help with figure preparation. We acknowledge generous support from the National Science Foundation CBET-1645123 (CMC and KS), EEC-1648035 (KS), AWD-101645-G3/RJ375-G3 (KS), and NSF Graduate Research Fellowship Program DGE-1747503, to K.P.M and N.J.P.); National Institutes of Health/National Cancer Institute R01 CA215461 (CMC), National Institute of General Medical Sciences R35 GM119644-01 (KS), National Institutes of Health grant 1S10OD025225-01 funding the Cytek Aurora Spectral Cytometer (Carbone Cancer Center), Biotechnology Training Program T32GM008349 (K.P.M); University of Wisconsin Carbone Cancer Center Support Grant P30 CA014520 Cell Based Immunotherapy Supplement (CMC and KS); St. Baldrick’s – Stand up to Cancer Pediatric Dream Team Translational Research Grant SU2C-AACR-DT-27-17 (CMC); American Cancer Society Research Scholar grant RSG-18-104-01-LIB (CMC); and the MACC Fund (CMC). Stand Up to Cancer is a division of the Entertainment Industry Foundation. Research grants are administered by the American Association for Cancer Research, the Scientific Partner of SU2C. The contents of this article do not necessarily reflect the views or policies of the Department of Health and Human Services, nor does mention of trade names, commercial products, or organizations imply endorsement by the US Government.

## Authors’ contributions

These authors contributed equally: K.P.M. and N.J.P. K.P.M. and L.A.S. designed and cloned the VFC-CAR and VFC-Ctrl plasmids. K.P.M. developed the two-step purification procedure for HDR templates, and K.P.M. and N.J.P. optimized electroporation protocols. K.P.M. and N.J.P. isolated and cultured T cells and performed virus-free transfections. K.P.M. and A.D. performed viral transductions. K.P.M. and B.R. performed and analyzed scRNA-seq experiments. K.P.M., M.S., M.F., L.A.S., and A.D. performed flow cytometry. N.J.P. and B.R. performed and analyzed NGS experiments. N.J.P and M.F. performed cytokine experiments and N.J.P. analyzed cytokine data. M.F. conducted *in vivo* experiments. K.P.M., K.Sh., and L.S. planned and performed western blotting experiments. K.P.M., N.J.P. and A.A. performed *in vitro* co-cultures. C.L. performed the CHANGE-seq assay. K.P.M. and N.J.P. wrote the manuscript with input from all authors. S.T., K.S. and C.M.C. supervised the research. K.P.M., N.J.P., M.F., L.A.S., M.S., K.Sh., L.S., C.L., A.D., A.A., and B.R. performed experiments and analyzed the data.

---

**Online Supplemental Figure S1.**
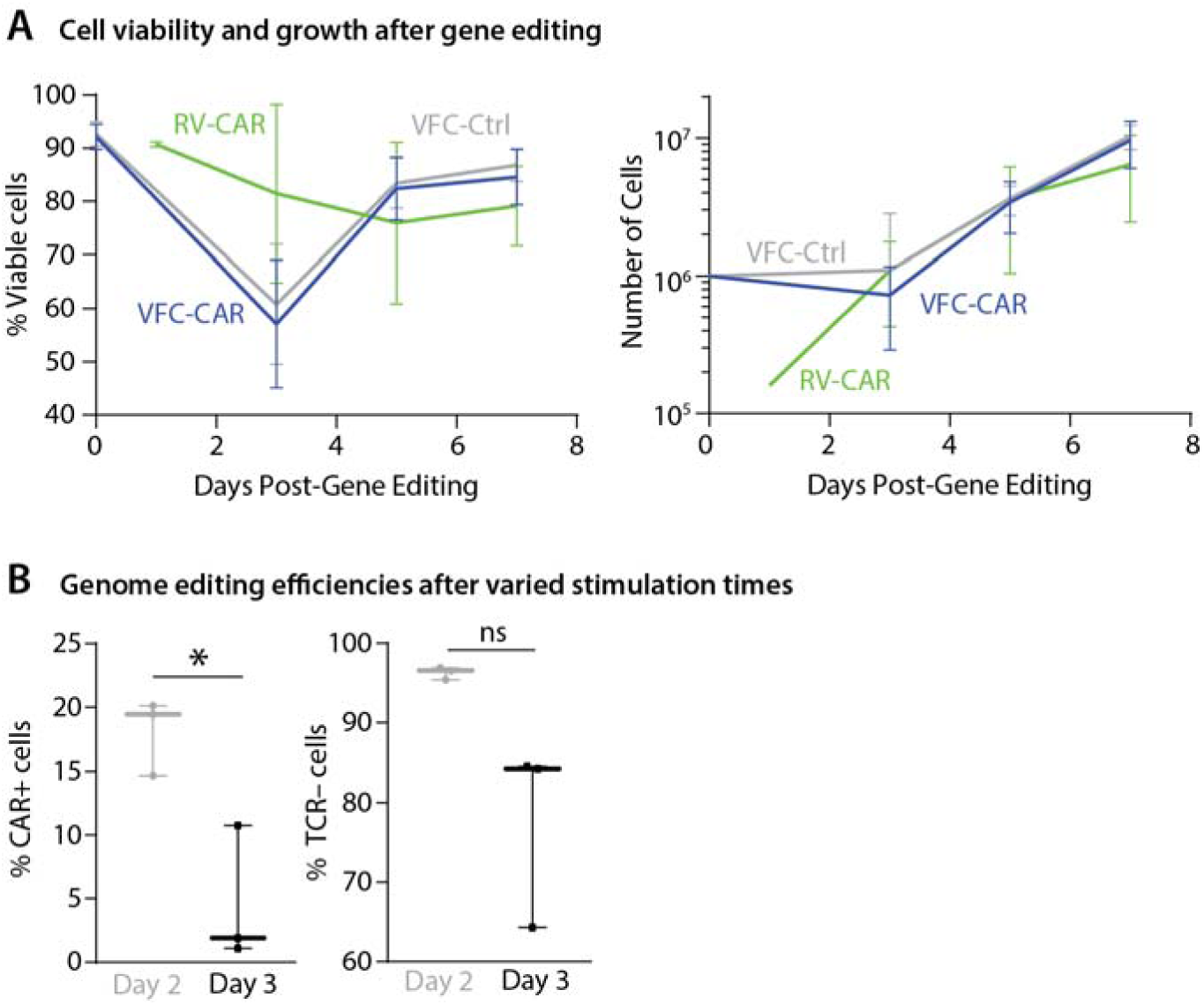
Characterization of T cell products during manufacturing. (A) Left, Viability of cells throughout the manufacturing timeline, pooled for 4 donors. Right, Cell counts throughout the manufacture calendar, pooled for 4 donors. VFC-CAR (blue) N=36; RV-CAR (green) N=27; VFC-Ctrl (gray) N=25. (B) Left, Percent of CAR+ cells as measured by flow cytometry when electroporated on day 2 or day 3 post-isolation. Right, Percent of TCR-cells as measured by flow cytometry when electroporated on day 2 or day 3 post-isolation. * indicates p<=0.05.

**Online Supplemental Figure S2.**
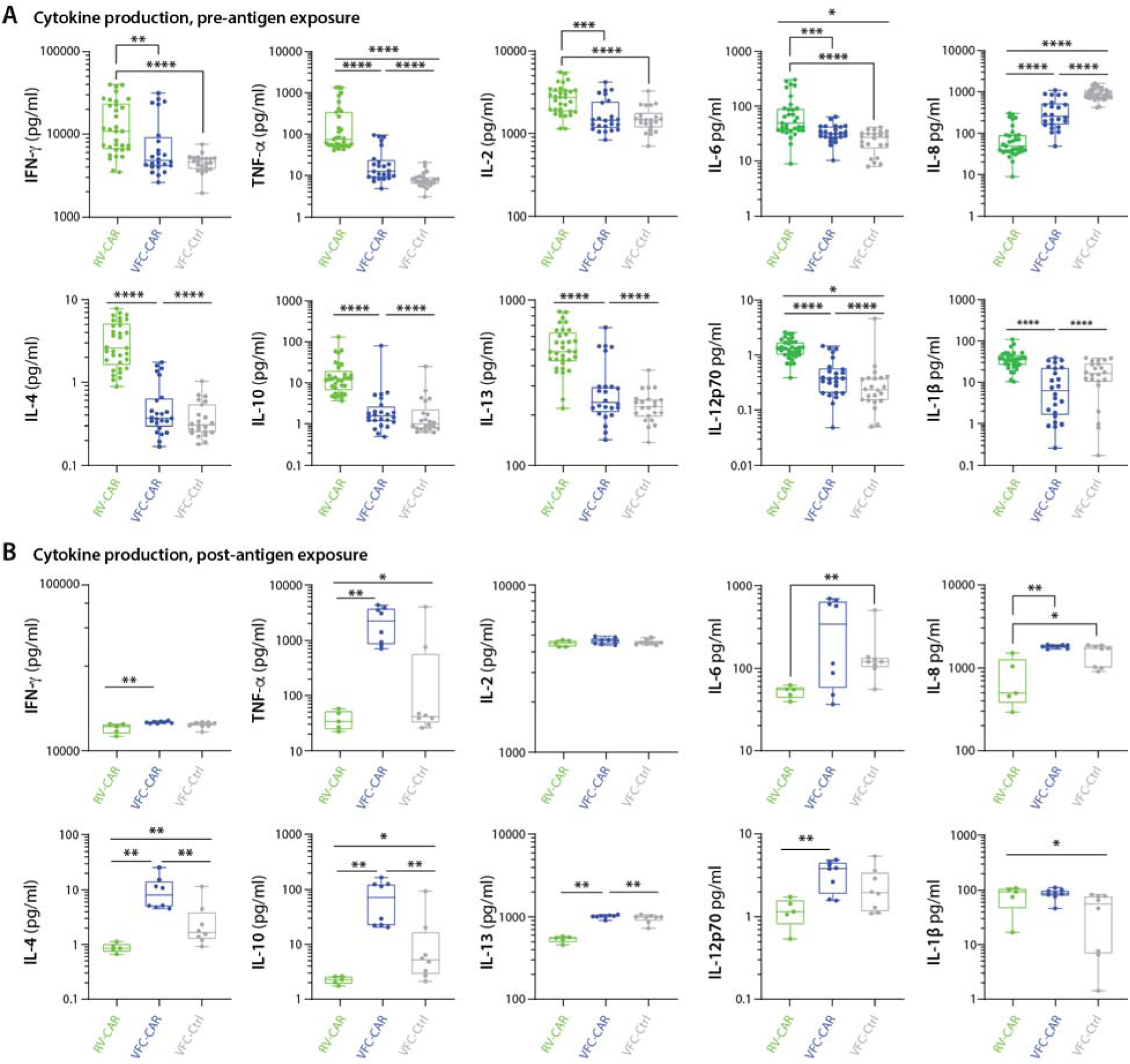
VFC-CAR T cells mount a robust cytokine response upon exposure to cognate antigen (individual replicates from figure 3A.) (A) Cytokine production from conditioned media taken from T cell products at the end of manufacturing (pre-antigen exposure). Values are pooled from four donors. VFC-CAR (blue) N=24; RV-CAR (green) N=33; VFC-Ctrl (gray) N=22. (B) Cytokine production in conditioned media after a 24 hour co-culture of manufactured T cell products with the target GD2-antigen on CHLA20 neuroblastoma cells. Values are pooled from two donors. VFC-CAR (blue) N=8; RV-CAR (green) N=5; VFC-Ctrl (gray) N=8. Statistical significance was calculated with a two-tailed Mann-Whitney test. * indicates p<=0.05; ** indicates p<=0.01; *** indicates p<=0.001; **** indicates p<=0.0001.

**Online Supplemental Figure S3.**
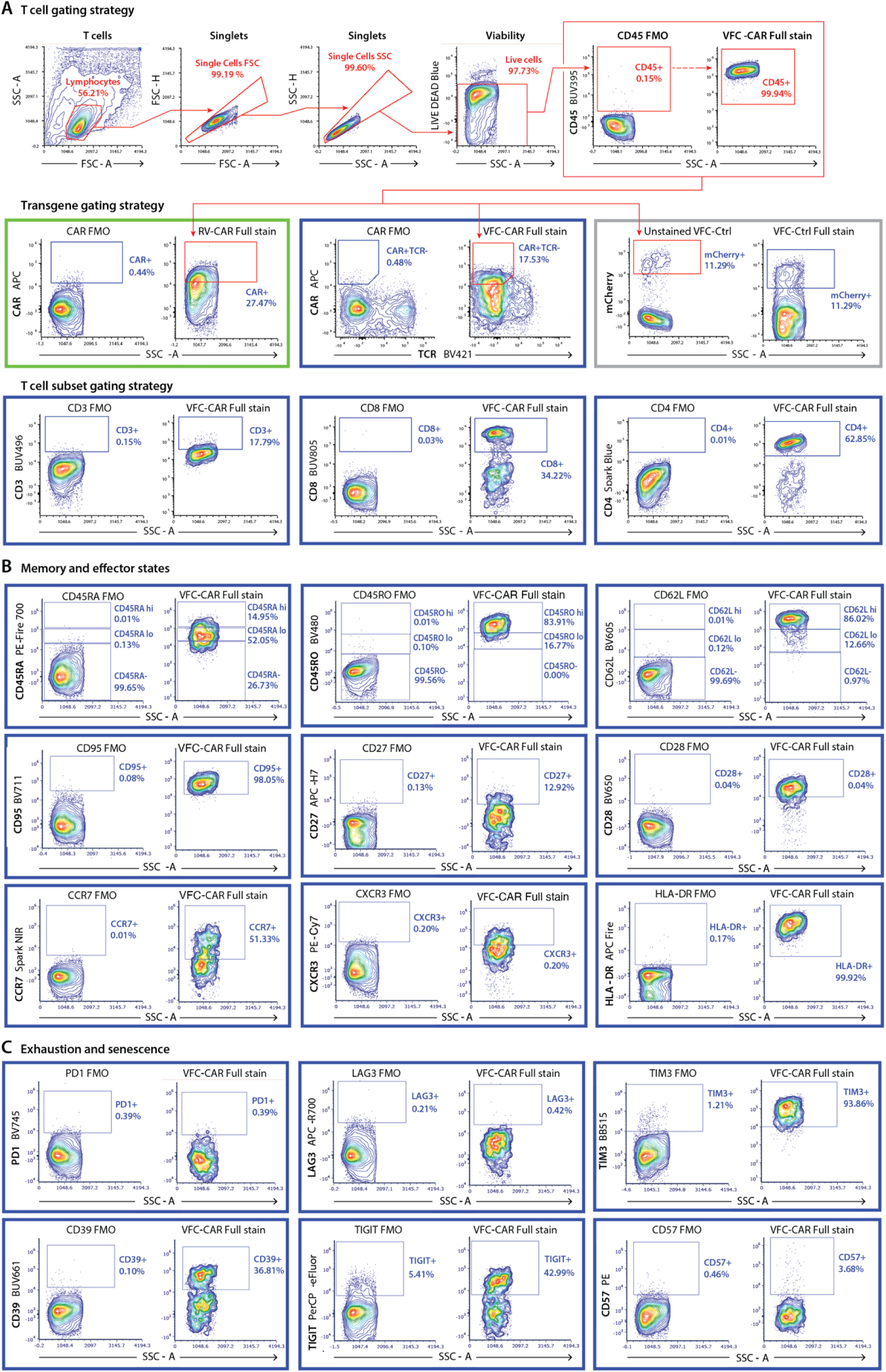
Immunophenotyping gating strategy. Cells were assayed by spectral cytometry with a 21-color immunophenotyping panel on day 10 of manufacture. (A) Cells were gated for lymphocytes, singlets, live cells, and CD45+ cells. (B) Subsequent gates for transgene+ cells were determined using fluorescence-minus-one (FMO) controls for each marker (left of each panel), with a representative full stain shown at right for each color. (C) Gating strategy for downstream immunophenotyping analysis of memory and effector states. Gates for each color were established using FMO controls, at left for each panel. All panels in blue boxes show representative data from one replicate of VFC-CAR T cells. Panel in green shows representative data from one replicate of RV-CAR T cells. Panel in grey shows representative data from one replicate of VFC-Ctrl T cells. All antibodies were titrated at 5 different concentrations to determine optimal staining conditions. SSC-A, side scatter area. FSC-A, forward scatter area. SSC-H, side scatter height. FSC-H, forward scatter height.

**Online Supplemental Figure S4.**
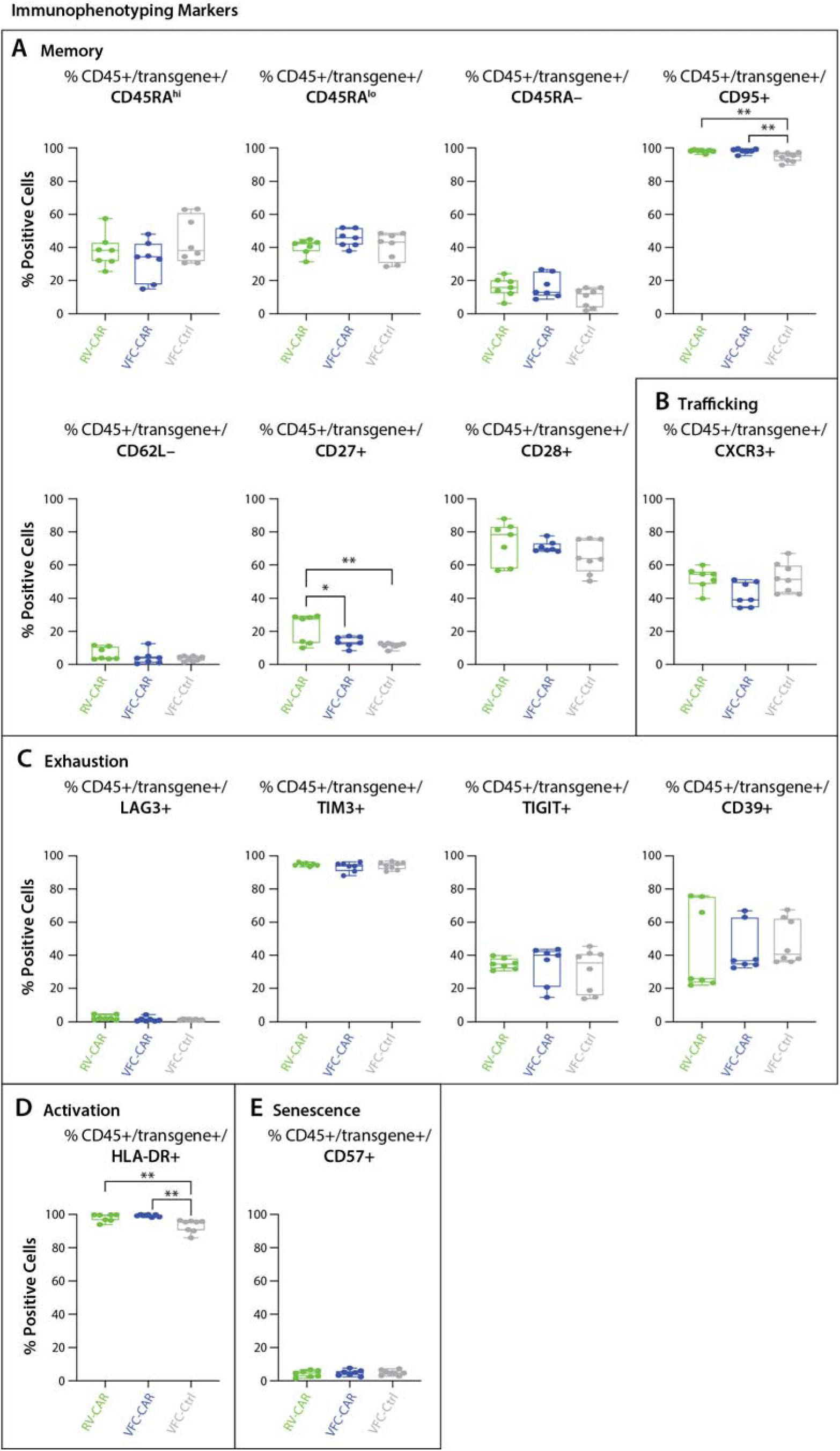
CAR T cell immunophnotypes. Cells were assayed by spectral cytometry with a 21-color immunophenotyping panel on day 10 of manufacture. (A) Additional markers are shown in figure 4. Gating strategies are shown in online supplemental **figure S3**. VFC-CAR (blue) N=7; RV-CAR (green) N=7; VFC-Ctrl (gray) N=8 across two donors. Significance was determined by ordinary one-way ANOVA; * indicates p≤0.05; **indicates p≤0.01; *** indicates p≤0.001; **** indicates p≤0.0001.

**Online Supplemental Figure S5.**
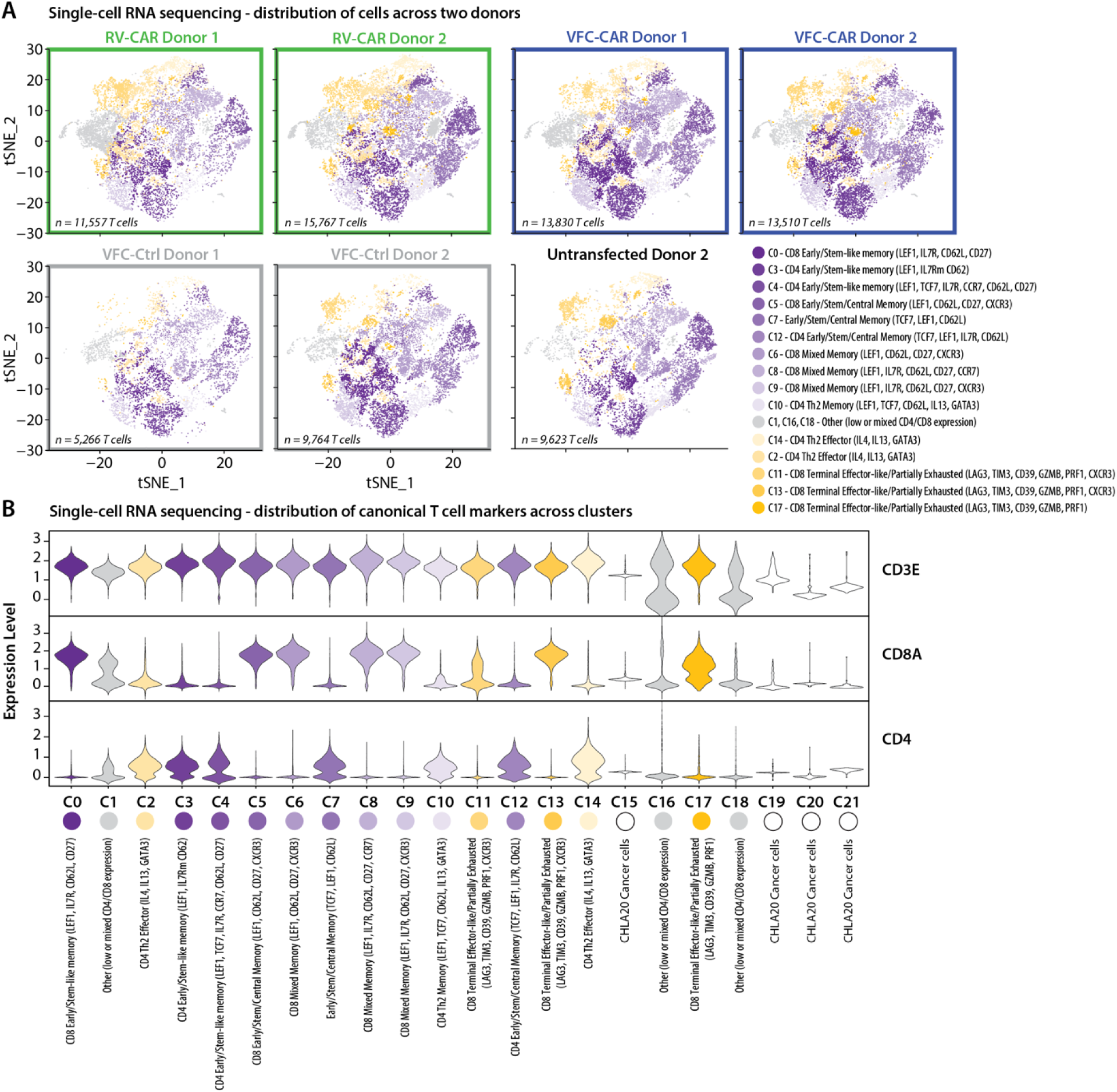
Transcriptional signatures of single CAR T cells. (A) tSNE projection of single cell RNA-seq data from 15 samples of manufactured cell products, both pre- and post-antigen exposure. Gross clustering patterns indicate similar cell populations across donors; each tSNE plot shows aggregate cells for each cell type both prior to and after antigen exposure, separated by donor. (B) Violin plots show expression of the pan-T cell markers CD3E, CD8A, and CD4. Clusters 15, 19, 20, and 21 showed decreased expressed of all three markers and were determined to be contaminating cancer cells from co-culture samples; these clusters are not shown on the tSNE plots and are excluded from all downstream analyses. Clusters 0, 5, 6, 8, 9, 13, and 17 were determined to contain CD8+ cytotoxic T cells; clusters 2, 3, 4, 7, 10, 12, and 14 were determined to contain CD4+ helper T cells. Clusters 1, 16 and 18 were determined to have mixed or low CD4/CD8 expression and were not annotated; they are designated as “other” in figure 5C and D. Expression level refers to log normalized data.

**Online Supplemental Figure S6.**
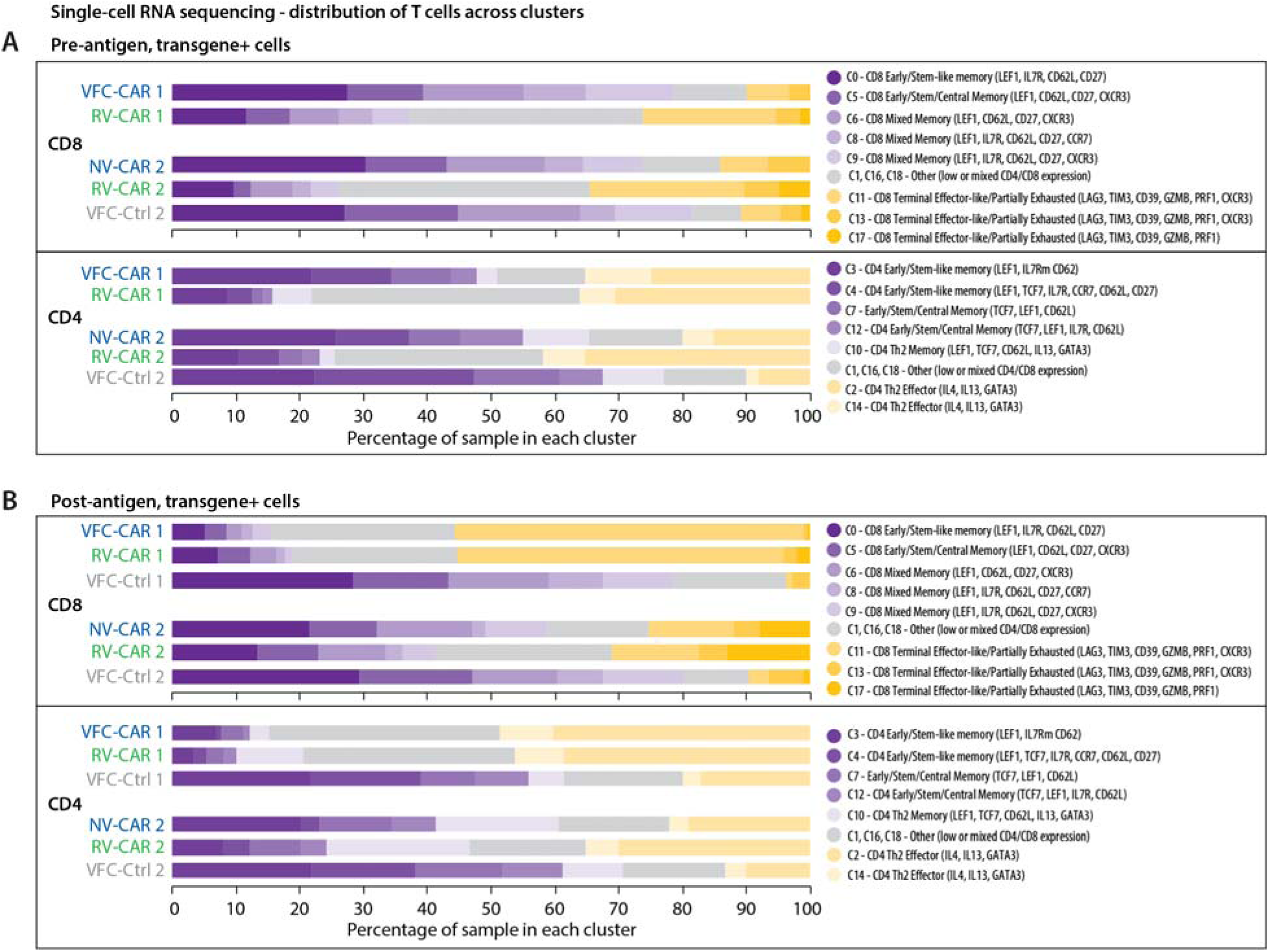
Transcriptional signatures of single CAR T cells prior to and after target antigen exposure. (A) Proportion of transgene+ cells from all pre-antigen samples within each annotated cluster. (B) Proportion of transgene+ cells from all post-antigen samples within each annotated cluster. Each color represents a different cluster, shown in figure 5A. Purple clusters are memory-associated; yellow clusters are effector-associated; grey clusters could not be identified as pure T cell clusters due to a mix or lack of robust CD4/CD8 expression. Bar charts are the same as those shown in figure 5C and D, separated by CD8 and CD4-specific clusters. Similar patterns of memory vs. effector formation were observed in both cytotoxic and helper T cells, with VFC-CAR and VFC-Ctrl cells skewing towards a memory phenotype prior to antigen exposure, and VFC-CAR and RV-CAR T cells acquiring an effector phenotype after antigen exposure.

**Online Supplemental Figure S7.**
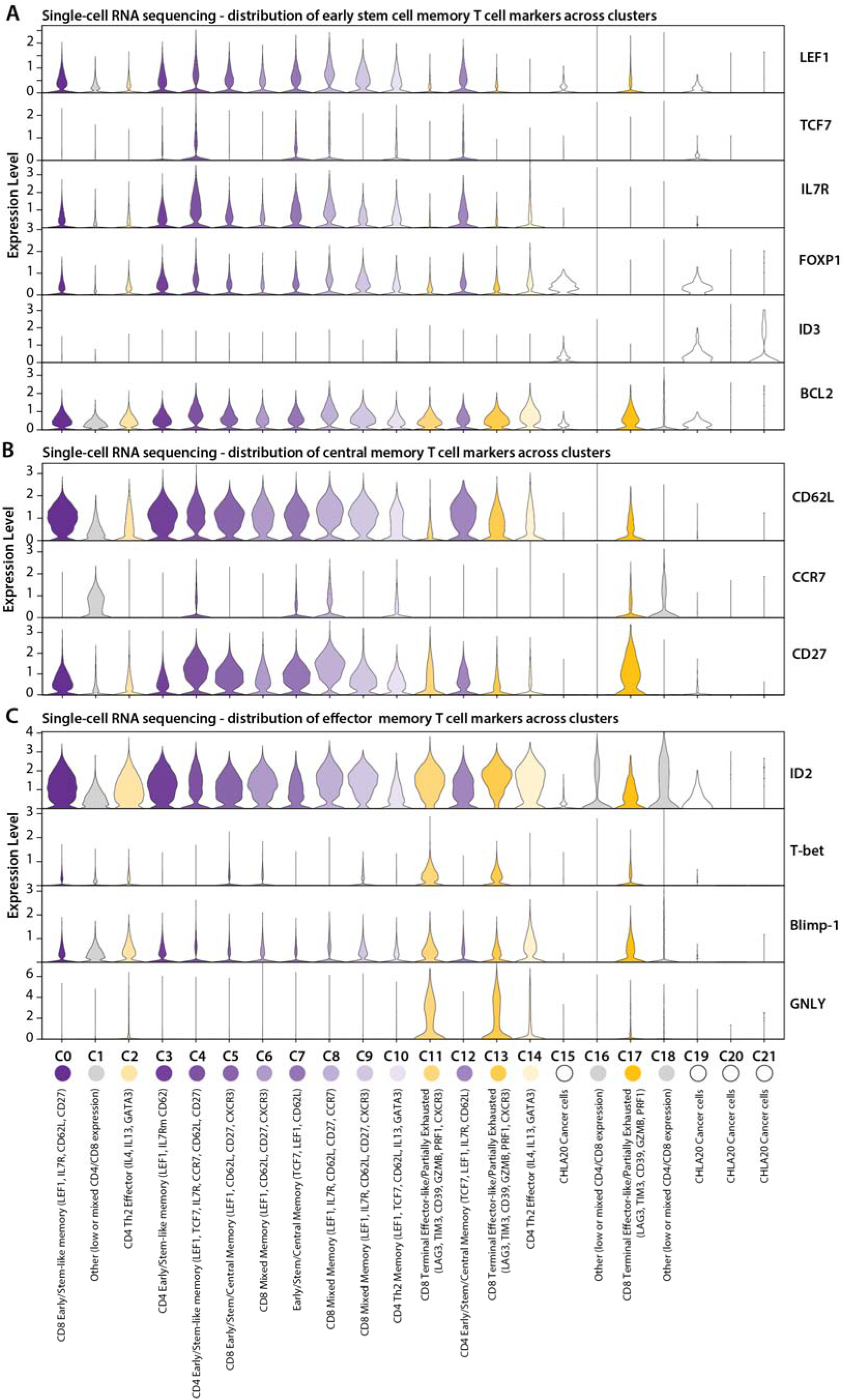
Memory-associated gene expression in single cell RNA-sequencing. (A) Violin plots show expression of six markers of early/stem cell memory phenotypes (LEF1, TCF7, IL7R, FOXP1, ID3, and BCL2). (B) Violin plots show expression of three markers of central memory T cells (CD62L, CCR7, and CD27). (C) Violin plots show expression of four markers of effector memory T cells (ID2, T-bet, Blimp-1, and GNLY). Clusters 0, 3, and 4 were determined to predominantly express early/stem-like memory markers. Clusters 5, 7, and 12 were determined to express a mix of stem-like and central memory markers. Clusters 6, 8, and 9 were determined to express a mix of stem-like, central, and effector memory markers. Cluster 10 was determined to express a predominantly Th2 memory phenotype. Clusters 2 and 14 were determined to express a Th2 effector phenotype. Clusters 11, 13, and 17 were determined to express an effector phenotype along with some exhaustion markers. Clusters 1, 16, and 18 showed low or mixed CD4/CD8 expression and were not annotated. Clusters 15, 19, 20 and 21 were determined to be cancer cells and were not included in downstream analyses. Expression level refers to log normalized data.

**Online Supplemental Figure S8.**
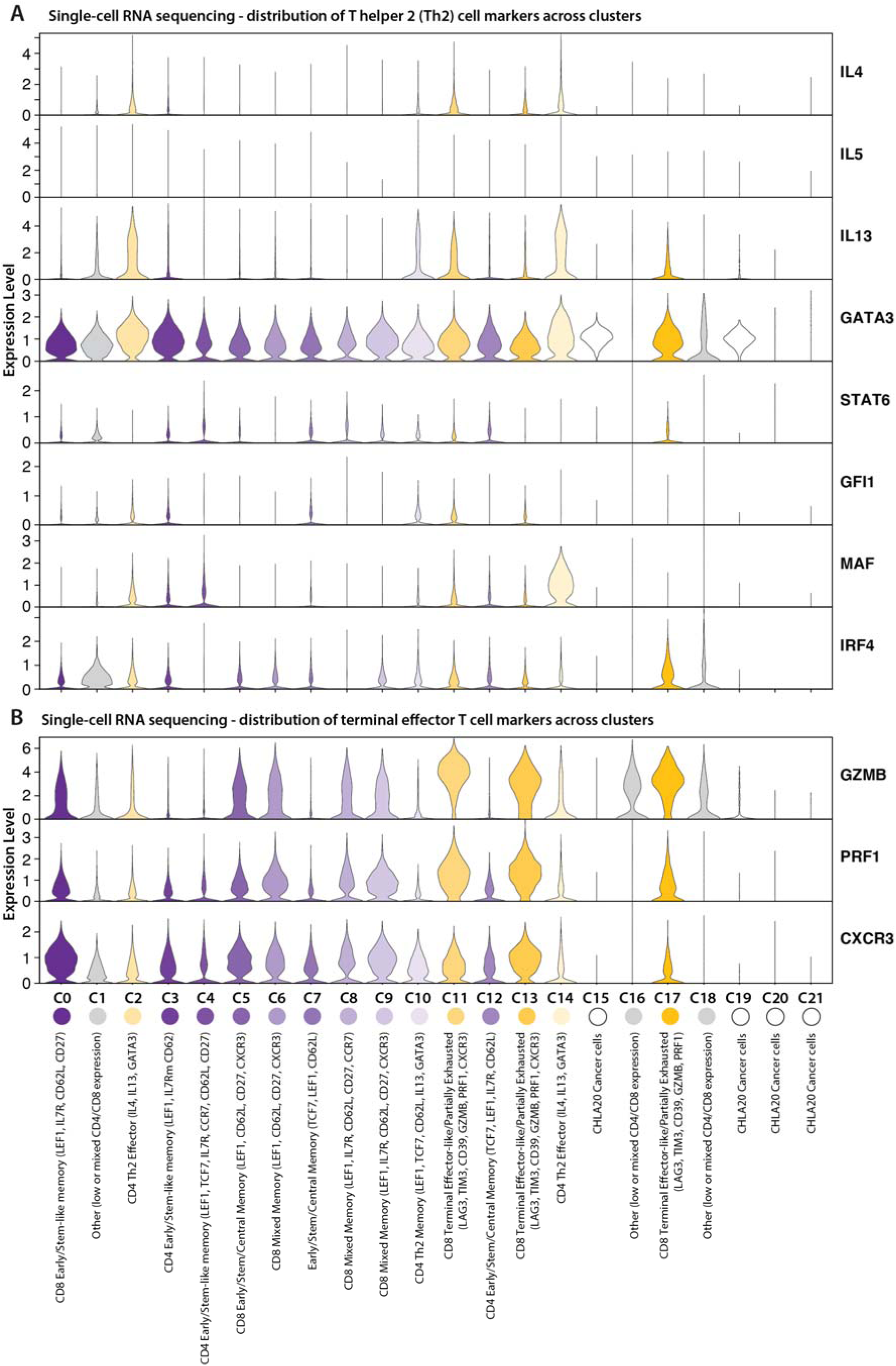
T helper 2 (Th2) and terminal effector gene expression in single cell RNA-sequencing. (A) Violin plots show expression of eight markers of Th2 helper T cells (IL4, IL5, IL13, GATA3, STAT6, GFI1, MAF, and IRF4). (B) (B) Violin plots show expression of three markers of terminal effector T cells (GZMB, PRF1, and CXCR3). Clusters 0, 3, and 4 were determined to predominantly express early/stem-like memory markers. Clusters 5, 7, and 12 were determined to express a mix of stem-like and central memory markers. Clusters 6, 8, and 9 were determined to express a mix of stem-like, central, and effector memory markers. Cluster 10 was determined to express a predominantly Th2 memory phenotype. Clusters 2 and 14 were determined to express a Th2 effector phenotype. Clusters 11, 13, and 17 were determined to express an effector phenotype along with some exhaustion markers. Clusters 1, 16, and 18 showed low or mixed CD4/CD8 expression and were not annotated. Clusters 15, 19, 20 and 21 were determined to be cancer cells and were not included in downstream analyses. Expression level refers to log normalized data.

**Online Supplemental Figure S9.**
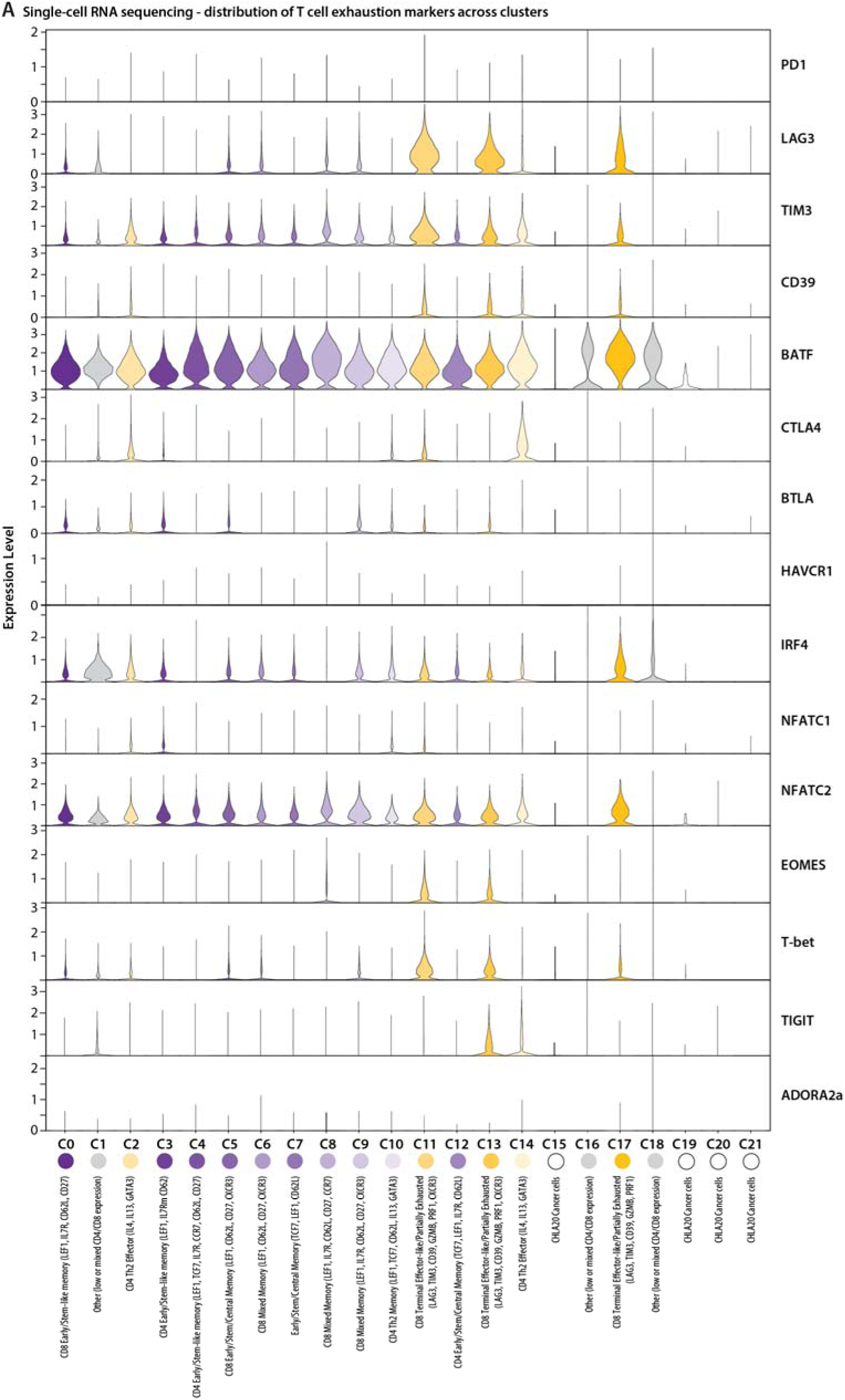
Exhaustion-associated gene expression in single cell RNA-sequencing. (A) Violin plots show expression of fifteen markers of exhaustion in T cells (PD1, LAG3, TIM3, CD39, BATF, CTLA4, BTLA, HAVCR1, IRF4, NFATC1, NFATC2, EOMES, T-bet, TIGIT, ADORA2A). Clusters 0, 3, and 4 were determined to predominantly express early/stem-like memory markers. Clusters 5, 7, and 12 were determined to express a mix of stem-like and central memory markers. Clusters 6, 8, and 9 were determined to express a mix of stem-like, central, and effector memory markers. Cluster 10 was determined to express a predominantly Th2 memory phenotype. Clusters 2 and 14 were determined to express a Th2 effector phenotype. Clusters 11, 13, and 17 were determined to express an effector phenotype along with some exhaustion markers. Clusters 1, 16, and 18 showed low or mixed CD4/CD8 expression and were not annotated. Clusters 15, 19, 20 and 21 were determined to be cancer cells and were not included in downstream analyses. Expression level refers to log normalized data.

**Online Supplemental Figure S10.**
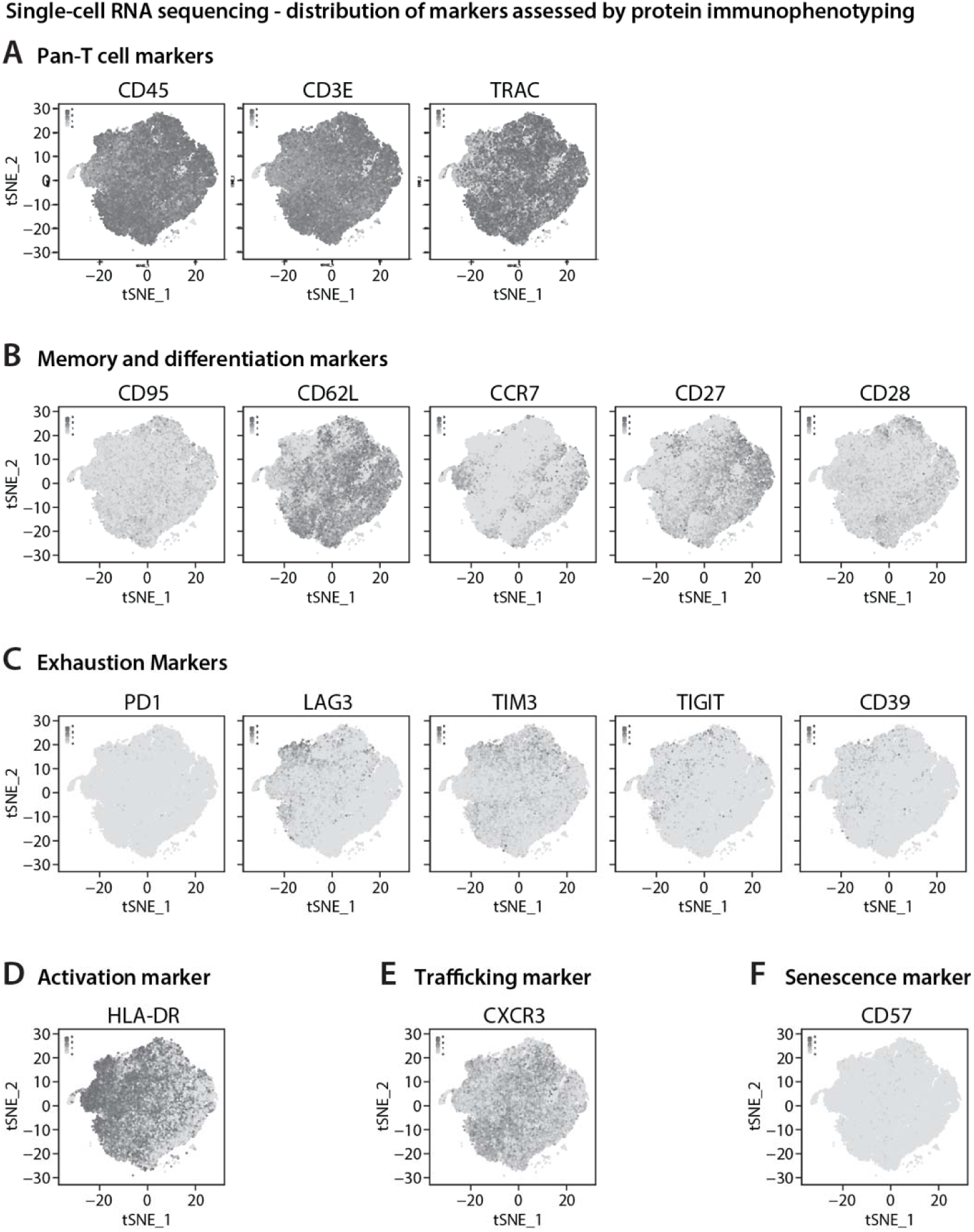
Single Cell RNA sequencing immunophenotyping. tSNE plots as shown in figure 5, colored for expression levels of all markers assayed for protein-level expression (figures 4 and S4). (A) Feature plots show expression of the pan T cell markers CD45, CD3E and TRAC. (B) Feature plots show expression of the memory and differentiation markers CD95, CD62L, CCR7, CD27, and CD28. (C) Feature plots show expression of the exhaustion markers PD1, LAG3, TIM3, TIGIT, and CD39. (D) Feature plot shows expression of the activation marker HLA-DR. (E) Feature plot shows expression of the T cell trafficking/inflammatory marker CXCR3. (F) Feature plot shows expression of the senescence marker CD57.

**Online Supplemental Figure S11.**
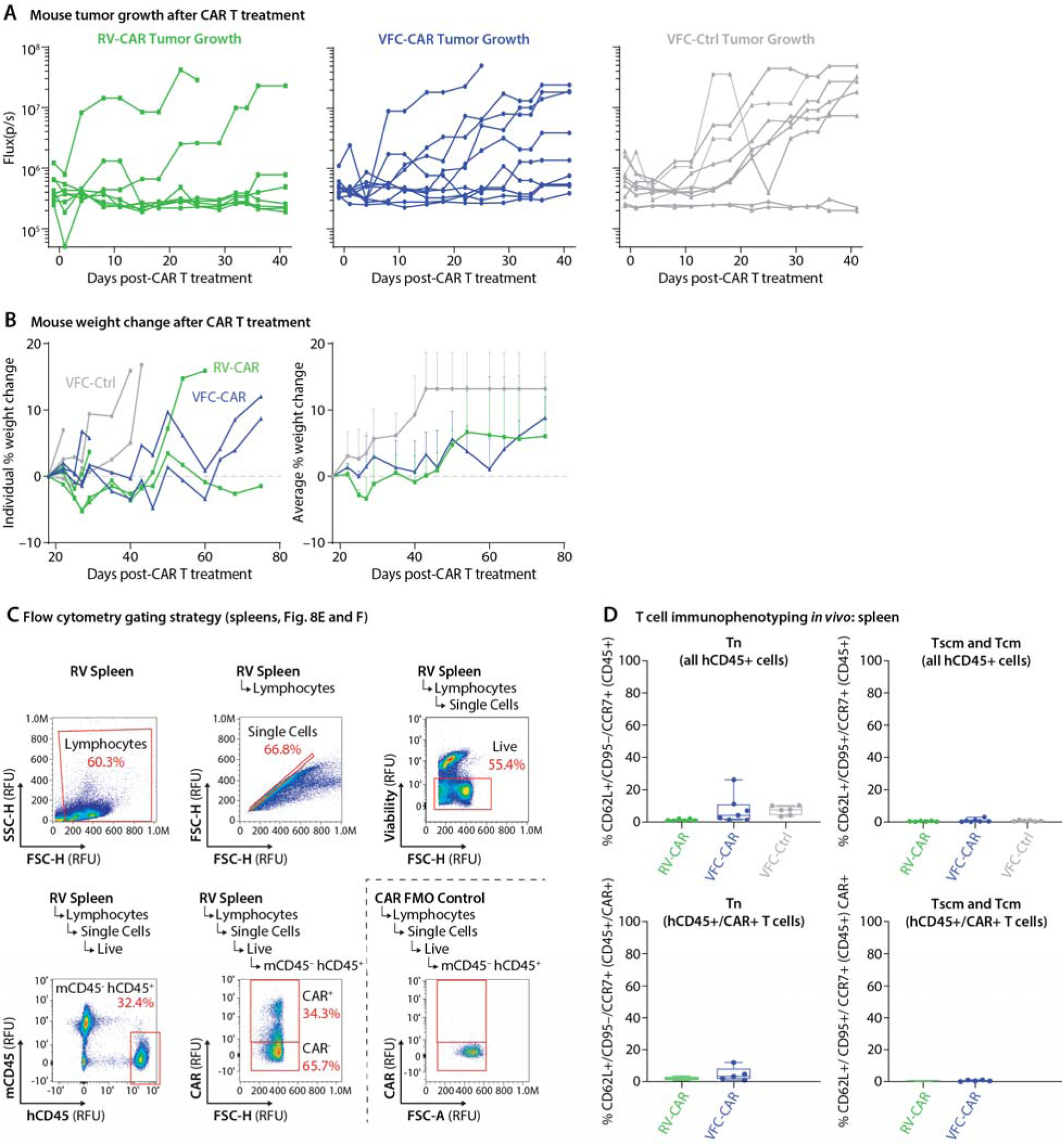
Bioluminescence, tumor growth, weight gains, T cell persistence, and memory formation after T cell treatment *in vivo*. (A) Flux measurements for individual luciferase-positive tumors for all mouse experiments. VFC-CAR, N=10. RV-CAR, N=8. VFC-Ctrl, N=7. (B) Left, individual mouse percent weight change throughout the experiment. Right, average percent weight change in mice per treatment condition. (C) Flow cytometric gating strategy used to assay mouse spleens for human T cells and CAR-positive cells. (D) Boxplots showing the expression levels of naïve (T_n_), stem cell memory (T_scm_) and central memory (T_cm_) markers on human T cells found in mouse spleens. For immunophenotyping: RV-CAR, N=6. VFC-Car, N=7. VFC-Ctrl, N=6.

## References

1. Goff, S. L. et al. Pilot Trial of Adoptive Transfer of Chimeric Antigen Receptor-transduced T Cells Targeting EGFRvIII in Patients With Glioblastoma. J. Immunother. 42, 126–135 (2019).

2. O’Rourke, D. M. et al. A single dose of peripherally infused EGFRvIII-directed CAR T cells mediates antigen loss and induces adaptive resistance in patients with recurrent glioblastoma. Sci. Transl. Med. 9, (2017).

3. Kahlon, K. S. et al. Specific recognition and killing of glioblastoma multiforme by interleukin 13-zetakine redirected cytolytic T cells. Cancer Res. 64, 9160–9166 (2004).

4. Newick, K., Moon, E. & Albelda, S. M. Chimeric antigen receptor T-cell therapy for solid tumors. Mol Ther Oncolytics 3, 16006 (2016).

5. Rodriguez, E., Schetters, S. T. T. & van Kooyk, Y. The tumour glyco-code as a novel immune checkpoint for immunotherapy. Nat. Rev. Immunol. 18, 204–211 (2018).

6. Fesnak, A. D., June, C. H. & Levine, B. L. Engineered T cells: the promise and challenges of cancer immunotherapy. Nat. Rev. Cancer 16, 566–581 (2016).

7. Piscopo, N. J. et al. Bioengineering Solutions for Manufacturing Challenges in CAR T Cells. Biotechnol. J. 13, (2018).

8. Wang, X. & Rivière, I. Clinical manufacturing of CAR T cells: foundation of a promising therapy. Mol Ther Oncolytics 3, 16015 (2016).

9. Nobles, C. L. et al. CD19-targeting CAR T cell immunotherapy outcomes correlate with genomic modification by vector integration. J. Clin. Invest. 130, 673–685 (2020).

10. van der Loo, J. C. M. & Wright, J. F. Progress and challenges in viral vector manufacturing. Hum. Mol. Genet. 25, R42–52 (2016).

11. Monjezi, R. et al. Enhanced CAR T-cell engineering using non-viral Sleeping Beauty transposition from minicircle vectors. Leukemia 31, 186–194 (2017).

12. Foster, J. B. et al. Purification of mRNA Encoding Chimeric Antigen Receptor Is Critical for Generation of a Robust T-Cell Response. Hum. Gene Ther. 30, 168–178 (2019).

13. Chong, E. A., Ruella, M., Schuster, S. J. & Lymphoma Program Investigators at the University of Pennsylvania. Five-Year Outcomes for Refractory B-Cell Lymphomas with CAR T-Cell Therapy. N. Engl. J. Med. 384, 673–674 (2021).

14. Jinek, M. et al. A programmable dual-RNA-guided DNA endonuclease in adaptive bacterial immunity. Science 337, 816–821 (2012).

15. McGuire, A. L. et al. The road ahead in genetics and genomics. Nat. Rev. Genet. 21, 581–596 (2020).

16. Doudna, J. A. The promise and challenge of therapeutic genome editing. Nature 578, 229–236 (2020).

17. Eyquem, J. et al. Targeting a CAR to the TRAC locus with CRISPR/Cas9 enhances tumour rejection. Nature 543, 113–117 (2017).

18. Sachdeva, M. et al. Repurposing endogenous immune pathways to tailor and control chimeric antigen receptor T cell functionality. Nat. Commun. 10, 5100 (2019).

19. Fraietta, J. A. et al. Determinants of response and resistance to CD19 chimeric antigen receptor (CAR) T cell therapy of chronic lymphocytic leukemia. Nat. Med. 24, 563–571 (2018).

20. Gardner, R. A. et al. Intent-to-treat leukemia remission by CD19 CAR T cells of defined formulation and dose in children and young adults. Blood 129, 3322–3331 (2017).

21. Feucht, J. et al. Calibration of CAR activation potential directs alternative T cell fates and therapeutic potency. Nat. Med. 25, 82–88 (2019).

22. Kalos, M. et al. T cells with chimeric antigen receptors have potent antitumor effects and can establish memory in patients with advanced leukemia. Sci. Transl. Med. 3, 95ra73 (2011).

23. Xu, Y. et al. Closely related T-memory stem cells correlate with in vivo expansion of CAR.CD19-T cells and are preserved by IL-7 and IL-15. Blood 123, 3750–3759 (2014).

24. Zah, E. et al. Systematically optimized BCMA/CS1 bispecific CAR-T cells robustly control heterogeneous multiple myeloma. Nat. Commun. 11, 2283 (2020).

25. Shah, N. N. & Fry, T. J. Mechanisms of resistance to CAR T cell therapy. Nat. Rev. Clin. Oncol. 16, 372–385 (2019).

26. Lynn, R. C. et al. c-Jun overexpression in CAR T cells induces exhaustion resistance. Nature 576, 293–300 (2019).

27. Louis, C. U. et al. Antitumor activity and long-term fate of chimeric antigen receptor-positive T cells in patients with neuroblastoma. Blood 118, 6050–6056 (2011).

28. Hanlon, K. S. et al. High levels of AAV vector integration into CRISPR-induced DNA breaks. Nat. Commun. 10, 4439 (2019).

29. Roth, T. L. et al. Reprogramming human T cell function and specificity with non-viral genome targeting. Nature 559, 405–409 (2018).

30. Kath, J. et al. Fast, efficient and virus-free generation of TRAC-replaced CAR T cells. bioRxiv 2021.02.14.431017 (2021) doi:10.1101/2021.02.14.431017.

31. Shy, B. R. et al. Hybrid ssDNA repair templates enable high yield genome engineering in primary cells for disease modeling and cell therapy manufacturing. doi:10.1101/2021.09.02.458799.

32. Roth, T. L. et al. Pooled Knockin Targeting for Genome Engineering of Cellular Immunotherapies. Cell 181, 728–744.e21 (2020).

33. Pulè, M. A. et al. A chimeric T cell antigen receptor that augments cytokine release and supports clonal expansion of primary human T cells. Mol. Ther. 12, 933–941 (2005).

34. Goodwin, M. et al. CRISPR-based gene editing enables FOXP3 gene repair in IPEX patient cells. Science Advances 6, (2020).

35. Lazzarotto, C. R. et al. CHANGE-seq reveals genetic and epigenetic effects on CRISPR-Cas9 genome-wide activity. Nat. Biotechnol. (2020) doi:10.1038/s41587-020-0555-7.

36. Rasaiyaah, J., Georgiadis, C., Preece, R., Mock, U. & Qasim, W. TCRαβ/CD3 disruption enables CD3-specific antileukemic T cell immunotherapy. JCI Insight 3, (2018).

37. Long, A. H. et al. 4-1BB costimulation ameliorates T cell exhaustion induced by tonic signaling of chimeric antigen receptors. Nat. Med. 21, 581–590 (2015).

38. Gomes-Silva, D. et al. Tonic 4-1BB Costimulation in Chimeric Antigen Receptors Impedes T Cell Survival and Is Vector-Dependent. Cell Rep. 21, 17–26 (2017).

39. Ajina, A. & Maher, J. Strategies to Address Chimeric Antigen Receptor Tonic Signaling. Mol. Cancer Ther. 17, 1795–1815 (2018).

40. Mahnke, Y. D., Brodie, T. M., Sallusto, F., Roederer, M. & Lugli, E. The who’s who of T-cell differentiation: human memory T-cell subsets. Eur. J. Immunol. 43, 2797–2809 (2013).

41. Summers, K. L., O’Donnell, J. L. & Hart, D. N. Co-expression of the CD45RA and CD45RO antigens on T lymphocytes in chronic arthritis. Clin. Exp. Immunol. 97, 39–44 (1994).

42. Li, Y. & Kurlander, R. J. Comparison of anti-CD3 and anti-CD28-coated beads with soluble anti-CD3 for expanding human T cells: differing impact on CD8 T cell phenotype and responsiveness to restimulation. J. Transl. Med. 8, 104 (2010).

43. Hao, Y. et al. Integrated analysis of multimodal single-cell data. Cell 184, 3573–3587.e29 (2021).

44. Sade-Feldman, M. et al. Defining T Cell States Associated with Response to Checkpoint Immunotherapy in Melanoma. Cell 175, 998–1013.e20 (2018).

45. Weber, E. W. et al. Transient rest restores functionality in exhausted CAR-T cells through epigenetic remodeling. Science 372, (2021).

46. Deng, Q. et al. Characteristics of anti-CD19 CAR T cell infusion products associated with efficacy and toxicity in patients with large B cell lymphomas. Nat. Med. (2020) doi:10.1038/s41591-020-1061-7.

47. Martin, M. D. & Badovinac, V. P. Defining Memory CD8 T Cell. Front. Immunol. 9, 2692 (2018).

48. Pule, M. A. et al. Virus-specific T cells engineered to coexpress tumor-specific receptors: persistence and antitumor activity in individuals with neuroblastoma. Nat. Med. 14, 1264–1270 (2008).

49. Heczey, A. et al. CAR T cells administered in combination with lymphodepletion and PD-1 inhibition to patients with neuroblastoma. Mol. Ther. 25, 2214–2224 (2017).

50. Hong, M., Clubb, J. D. & Chen, Y. Y. Engineering CAR-T Cells for Next-Generation Cancer Therapy. Cancer Cell (2020) doi:10.1016/j.ccell.2020.07.005.

51. Martinez, M. & Moon, E. K. CAR T Cells for Solid Tumors: New Strategies for Finding, Infiltrating, and Surviving in the Tumor Microenvironment. Front. Immunol. 10, 128 (2019).

52. Tokarew, N., Ogonek, J., Endres, S., von Bergwelt-Baildon, M. & Kobold, S. Teaching an old dog new tricks: next-generation CAR T cells. Br. J. Cancer 120, 26–37 (2019).

53. Tanna, J. G., Ulrey, R., Williams, K. M. & Hanley, P. J. Critical testing and parameters for consideration when manufacturing and evaluating tumor-associated antigen-specific T cells. Cytotherapy 21, 278–288 (2019).

54. U.S. Department of Health and Human Services FDA, Center for Biologics Evaluation and Research. Chemistry, Manufacturing, and Control (CMC) Information for Human Gene Therapy Investigational New Drug Applications (INDs) Guidance for Industry. https://www.fda.gov/regulatory-information/search-fda-guidance-documents/chemistry-manufacturing-and-control-cmc-information-human-gene-therapy-investigational-new-drug (01/31/2020).

55. Shin, J. J. et al. Controlled Cycling and Quiescence Enables Efficient HDR in Engraftment-Enriched Adult Hematopoietic Stem and Progenitor Cells. Cell Rep. 32, 108093 (2020).

56. Nguyen, D. N. et al. Polymer-stabilized Cas9 nanoparticles and modified repair templates increase genome editing efficiency. Nat. Biotechnol. 38, 44–49 (2020).

57. Kunz, A. et al. Optimized Assessment of qPCR-Based Vector Copy Numbers as a Safety Parameter for GMP-Grade CAR T Cells and Monitoring of Frequency in Patients. Mol Ther Methods Clin Dev 17, 448–454 (2020).

58. Lu, A. et al. Application of droplet digital PCR for the detection of vector copy number in clinical CAR/TCR T cell products. J. Transl. Med. 18, 191 (2020).

59. Nelson, C. E. et al. Long-term evaluation of AAV-CRISPR genome editing for Duchenne muscular dystrophy. Nat. Med. 25, 427–432 (2019).

60. Webber, B. R., et al. Highly efficient multiplex human T cell engineering without double-strand breaks using Cas9 base editors. Nat. Commun. 10, 5222 (2019).

61. Dai, X. et al. One-step generation of modular CAR-T cells with AAV-Cpf1. Nat. Methods 16, 247–254 (2019).

62. Ren, J. et al. A versatile system for rapid multiplex genome-edited CAR T cell generation. Oncotarget 8, 17002–17011 (2017).

63. Khajanchi, N. & Saha, K. Controlling CRISPR with Small Molecule Regulation for Somatic Cell Genome Editing. Mol. Ther. (2021) doi:10.1016/j.ymthe.2021.06.014.

64. Wu, L., Wei, Q., Brzostek, J. & Gascoigne, N. R. J. Signaling from T cell receptors (TCRs) and chimeric antigen receptors (CARs) on T cells. Cell. Mol. Immunol. 17, 600–612 (2020).

65. Maeda, T. et al. Regeneration of CD8αβ T Cells from T-cell-Derived iPSC Imparts Potent Tumor Antigen-Specific Cytotoxicity. Cancer Res. 76, 6839–6850 (2016).

66. Nagano, S. et al. High Frequency Production of T Cell-Derived iPSC Clones Capable of Generating Potent Cytotoxic T Cells. Mol Ther Methods Clin Dev 16, 126–135 (2020).

67. Themeli, M. et al. Generation of tumor-targeted human T lymphocytes from induced pluripotent stem cells for cancer therapy. Nat. Biotechnol. 31, 928–933 (2013).

68. Shukla, S. et al. Progenitor T-cell differentiation from hematopoietic stem cells using Delta-like-4 and VCAM-1. Nat. Methods 14, 531–538 (2017).

69. Liu, E. et al. Cord blood NK cells engineered to express IL-15 and a CD19-targeted CAR show long-term persistence and potent antitumor activity. Leukemia 32, 520–531 (2018).

70. Terakura, S. et al. Generation of CD19-chimeric antigen receptor modified CD8+ T cells derived from virus-specific central memory T cells. Blood 119, 72–82 (2012).

71. Cromer, M. K. et al. Global Transcriptional Response to CRISPR/Cas9-AAV6-Based Genome Editing in CD34+ Hematopoietic Stem and Progenitor Cells. Mol. Ther. 26, 2431–2442 (2018).

72. Viaud, S. et al. Switchable control over in vivo CAR T expansion, B cell depletion, and induction of memory. Proc. Natl. Acad. Sci. U. S. A. 115, E10898–E10906 (2018).

73. Hwang, J.-R., Byeon, Y., Kim, D. & Park, S.-G. Recent insights of T cell receptor-mediated signaling pathways for T cell activation and development. Exp. Mol. Med. 52, 750–761 (2020).

74. Wang, K. et al. A multiscale simulation framework for the manufacturing facility and supply chain of autologous cell therapies. Cytotherapy 21, 1081–1093 (2019).

75. Lesueur, L. L., Mir, L. M. & André, F. M. Overcoming the Specific Toxicity of Large Plasmids Electrotransfer in Primary Cells In Vitro. Mol. Ther. Nucleic Acids 5, e291 (2016).

76. Ruella, M. et al. Induction of resistance to chimeric antigen receptor T cell therapy by transduction of a single leukemic B cell. Nat. Med. 24, 1499–1503 (2018).

77. Clement, K., Hsu, J. Y., Canver, M. C., Joung, J. K. & Pinello, L. Technologies and Computational Analysis Strategies for CRISPR Applications. Mol. Cell 79, 11–29 (2020).

78. Vakulskas, C. A. et al. A high-fidelity Cas9 mutant delivered as a ribonucleoprotein complex enables efficient gene editing in human hematopoietic stem and progenitor cells. Nat. Med. 24, 1216–1224 (2018).

